# Differential contribution of elastin and fibrillin-1 to the cardiovascular phenotype of a double heterozygous Marfan and Williams-Beuren syndrome mouse model

**DOI:** 10.1101/2025.07.01.662517

**Authors:** Isaac Rodríguez-Rovira, Lize Sels, Jana Ruíz-Castro, Alba Aizpuru-Gómez, Ana Paula Dantas, Victoria Campuzano, Gustavo Egea

## Abstract

Marfan syndrome (MFS) and Williams-Beuren syndrome (WBS) are two genetic diseases of connective tissue caused respectively by mutations in the fibrillin-1 gene (FBN1) and hemizygous loss of the elastin gene (ELN) from an allelic chromosomic deletion. Their respective vascular manifestations are opposed, resulting in thoracic aortic aneurysm in MFS and supravalvular aortic stenosis in WBS. To investigate the interdependence of both essential molecular constituents of elastic fibers, we have generated a double heterozygous mouse model by crossing an MFS female (Fbn1C1041G/+) with a Complete Deletion (CD)-WBS male. We evaluated blood pressure, cardiac and aortic phenotypes, and some physical and cognitive functions in the offspring. Double heterozygous mice (CDMFS) presented the characteristic CD altered behavior. CDMFS mice developed an aneurysm, which progressed over age indistinguishably from MFS mice; CDMFS aortic wall showed a thicker tunica media aorta layer and thinner elastic fibers in accordance with CD mice. The characteristic MFS reduction in vascular smooth muscle cell density is not observed in the CDMFS aorta. CDMFS mice develop the high blood pressure observed in CD animals which does not occur in MFS. Ejection fraction was significantly reduced in MFS, CD, and CDMFS mice compared with WT littermates. Cardiac redox stress markers increased only in CD mice, but not in CDMFS animals. Metalloproteinase-2 protein levels increased only in MFS hearts and partially reduced in CDMFS ones. In conclusion, CDMFS mice exhibited the characteristic phenotypic manifestations of each syndrome, along with the pathological outcomes associated with each elastic fiber component and the respective disease.

## 1. INTRODUCTION

Elastic fibres are the second most abundant fibrous component of connective tissue after collagen fibres. They are basically composed of elastin and fibrillin-1 accompanied by other essential components such as fibulins, which provide the necessary, correct assembly of both basic components as well as the required biophysical and biochemical properties to be fully functionally integrated into the extracellular matrix ^1–3^. Elastic fibres can be histologically organised as individualised fibres and thin laminae and localised to numerous tissues and organs; when the elastic component is significantly present, these are grouped as elastic tissues, elastic arteries being the most enriched structures of the cardiovascular system. Of note, unlike elastin, fibrillin-1 has been reported to also be present in the heart ^4,5^. Two rare genetic diseases perturb elastic fibres, namely Marfan syndrome (MFS; OMIM 154700) and Williams-Beuren syndrome (WBS; OMIM 194050), which affect fibrillin-1 and elastin, respectively. Interestingly, despite both fibrillin-1 and elastin being the bases of elastic fibres, their pathological impact on the vascular system is opposed—aortic aneurysm in MFS and aortic stenosis in WBS ^6^.

MFS is caused by genetic variants in the fibrillin-1 gene (*FBN1*). Clinical characterisation is clearly defined by Ghent nosology by considering alterations in the cardiovascular (aortic root (AR) dilatation), muscle-skeleton (abnormally long bones and fingers), ocular (dislocation of the crystalline lens,) and respiratory (emphysema and sleep apnoea) systems ^7,8^. The usual disease progression of the aneurysm is aortic dissection, which usually ends with a fatal rupture of the aortic wall. Unfortunately, there is no clearly efficient pharmacological therapy, although beta-blockers and angiotensin II receptor blockers are clinically prescribed to slow down aneurysm dilatation ^9–12^. In the end, reparatory aortic surgery is currently the most efficient (and transient) solution ^13^. Several mouse models of the disease have been generated over time ^14^, which have provided definitive insights into our knowledge of disease pathogenesis and evolution. Thus, MFS pathomechanisms involve several dysfunctional interdependent molecular components and processes such as TGF-β and BMP signalling, extracellular matrix haemodynamic blood flow, redox stress, and endothelial and vascular smooth muscle cell (VSMC) disarrays, which together result in severe abnormal mechanobiology of the aortic wall ^15^.

On the other hand, WBS presents stenotic lesions in the thoracic aorta, although pulmonary and renal arteries are also altered ^16^. High blood pressure and supra-valvular ascending aortic stenosis (SVAS) are the most common cardiovascular manifestations of WBS ^16^. The arteriopathy is associated with VSMC hyperplasia and thinner elastic fibres, accompanied by an increased number of elastic lamellae ^17^. Severe SVAS usually leads to cardiac hypertrophy, increasing the risk of complications such as stroke and sudden death ^16,18^. These cardiovascular features are mainly related to a gene-dosage effect due to hemizygosity of the elastin gene (*ELN*), which encodes tropoelastin. Nonetheless, the phenotypic variability among WBS patients suggests the involvement of significant additional genetic modifiers that contribute to the clinical presentation beyond the effects of elastin deficiency alone ^17^. In our lab, we have generated a mouse model of the disease (CD model) that carries the most common deletion found in WBS patients (from *Gtf2i* to *Fkbp6*). This mouse model displays the characteristic pathological cardiovascular phenotype (SVAS, hypertrophic cardiopathy, and hypertension), being highly similar to that seen in patients ^19,20^. Moreover, the CD mouse model is the most suitable for assaying potential new therapeutic interventions ^21–24^.

The present study aimed to dissect the potential interdependence of elastin and fibrillin-1 genetic alterations in the appearance of the characteristic cardiovascular manifestations in a double heterozygous mouse model (CDMFS) generated from the well-characterised CD model of WBS ^19^ and MFS *Fbn1^C1041G/+^* ^25^. In addition to the analysis of a wide variety of cardiovascular outcomes, we also evaluated natural behaviour with a very simple test since cognitive dysfunctions are also characteristic of patients with WBS ^26^ and are present in the CD model ^27,28^. Our study complements and deepens a previous study where both elastin and fibrillin-1 were haploinsufficient and that mainly focused on mechanical vascular properties ^29^. Our study confirms some of the vascular findings observed in the haploinsufficiency *Eln^+/-^-Fbn1^+/-^*mouse model, such as aortic wall disarrays ^29^. However, our novel MFS-WBS combined mouse model (CDMFS) addresses new aortic and cardiac insights, such as aortic histopathology and transmission electron microscopy, aortic media proliferation/apoptosis, redox stress, mitochondrial morphology, and respiratory chain composition. In addition, we have also investigated basic brain and behaviour alterations specific to WBS.

## 2. MATERIAL AND METHODS

### 2.1 Ethics

All animal procedures were conducted following ARRIVE (Animals in Research: Reporting In Vivo Experiments) guidelines ^30^ and standard ethical guidelines (European Union Directive 2010/63/EU) and were approved by the local ethics committee (Animal Experimentation Ethics Committee-University of Barcelona, CEEA-UB) and the Government of Catalonia.

### 2.2 Animal models

Mice harbouring a fibrillin-1 mutation (*Fbn1^C1041G/+^*, referred to as C1041G and previously as C1039G/+; hereafter MFS mice) were obtained from The Jackson Laboratory (B6.129-Fbn1tm1Hcd/J; Strain #012885). CD mice were obtained as previously described ^19^. All mice were maintained in a C57BL/6J background (backcrossed for nine generations). We obtained the CD-Marfan mice colony (CDMFS) from the breeding of MFS females with CD males. The opposite breeding of MFS males with CD females was discarded because CD females are known to give birth to poor litters, which, moreover, they neglect ^19^. The experimental groups were wild-type (WT), MFS (Marfan syndrome), CD (WBS), and CDMFS (double heterozygous CD/MFS) mice. Mice were housed in controlled environmental conditions (21 ± 1°C temperature and 55 ± 10% humidity), and food and water were available *ad libitum*. All experiments were performed during the light phase of a 12-h light/dark cycle (light on at 8 am; light off at 8 pm). Genomic DNA was extracted from mouse ear punch to determine the genotype of each mouse using PCR, as previously described ^23^.

### 2.3 Marble Burying Test

The Marble Burying Test (MBT) was conducted in a polycarbonate rat cage filled with bedding to a depth of 5 cm and lightly tamped down. Twenty evenly spaced glass marbles (five rows of four marbles) were placed on the surface of the bedding before each test. One animal was placed in each cage. The number of buried marbles (> 2/3 of the marble covered) was counted after 20 minutes. In all experiments, investigators were blinded to assayed genotypes.

### 2.4 Blood pressure measurement

Systolic blood pressure (SBP) was measured in conscious mice using a tail-cuff system plethysmometer (Non-Invasive Blood Pressure System, PanLab, Barcelona, Spain) while the mice were placed in a restrainer tube, which was cleaned after each use. Following habituation to the restrainer, measurements were taken on three separate occasions and averaged for each mouse.

### 2.5 Echocardiography

Two-dimensional transthoracic echocardiography was performed on all animals under 1.5% inhaled isoflurane. Imaging was conducted using a 10–13 MHz phased array linear transducer (IL12i, GE Healthcare, Madrid, Spain) with a Vivid Q system (GE Healthcare, Madrid, Spain). The images were recorded and later analysed offline using EchoPac software (v.08.1.6, GE Healthcare, Madrid, Spain). Proximal aortic segments were examined in a parasternal long-axis view, and the AR diameter was measured from inner edge to inner edge at end-diastole at the level of the sinus of Valsalva. The left ventricular end-diastolic (LVDD) and end-systolic (LVSD) internal diameters were measured in M-mode recordings and computed by the Teichholz formula ^31^. The interventricular septum (IVSd) and left ventricular posterior wall thickness (LVWPd) in end diastole were also recorded. The average of three consecutive cardiac cycles was used for each measurement, with the operator being blinded to group assignment.

### 2.6 Tissue histopathology and electron microscopy

Paraffin-embedded tissue arrays (TMAs) of mice aortae and hearts from all experimental groups were transversally sectioned into 5-µm slices. Heart sections were stained with a regular haematoxylin-eosin protocol. Aorta sections were stained with Verhoeff–Van Gieson (Merck HT25A) following the supplier’s recommendations. Images were obtained under visible light using a Leica Leitz DMRB microscope (10X or 20X) equipped with a Leica DC500 camera. All measurements were carried out blindly, with no knowledge of genotypes.

Elastic fibre ruptures were quantified by counting breaks > 20 µm (large discontinuities) in the normal 360° circumferential continuity of each elastic lamina in the aortic media. These breaks were counted in four representative images from three non-consecutive sections of the same ascending aorta.

The undulation factor is defined as the ratio of elastic fibre size to the number of contact points where a continuous elastic fibre intersects or touches a virtually established straight line (value “=1” means that the fibre is completely straight; value “>1” indicates the undulation level of the fibre; the higher value, the greater the undulation).

For VSMC density and elastic fibre autofluorescence analysis, aortic sections were deparaffinised and rehydrated before DAPI staining and mounted on the same glass slide using a fluorescence-compatible mounting medium. Images were obtained using a Widefield Leica AF6000 (40X oil immersion) equipped with a Leica DC500 camera. For electron microscopy, isolated ascending aortae from mice of the four experimental groups were cut into rings and fixed with 2.5% glutaraldehyde and 2% paraformaldehyde in 0.1 M phosphate buffer, postfixed with 1% OsO_4_, dehydrated with ethanol, properly oriented, and embedded in Spurr resin before sectioning with a Leica ultramicrotome UC7 (Leica Microsystems). Ultrathin sections were not counterstained and were directly observed under a JEOL JEM-1010 TEM fitted with a Gatan Orius SC1000 (model 832) digital camera.

### 2.7 Immunohistochemistry

Paraffin-embedded cardiac tissue sections were deparaffinised and rehydrated before epitope unmasking when required. For this, sections were treated with a retrieval solution (10 mM sodium citrate, 0.05% Tween, pH 6) for 30 min in a steamer at 95°C. Sections were then incubated with ammonium chloride (NH_4_Cl, 50 mM, pH 7.4) for 20 min to block free aldehyde groups, followed by permeabilization with 0.3% Triton X-100 for 10 min. After rinsing three times with phosphate-buffered saline (PBS), sections were incubated with 1% bovine serum albumin (BSA) in PBS for 2 h at room temperature and then placed in a humidified chamber overnight at 4°C with the primary antibody (anti-3-NT, 1:200; Merck Millipore 06-284). The next day, sections were incubated with an anti-rabbit secondary antibody-Alexa 555 (1:1000, Invitrogen) at room temperature for 60 min. Finally, sections were stained with DAPI and mounted on the same glass slide using a fluorescence-compatible mounting medium. Images were obtained using a Widefield Leica AF6000 (40X oil immersion) equipped with a Leica DC500 camera.

### 2.8 In-situ detection of superoxide anions

Deparaffinised and rehydrated paraffin-embedded cardiac sections were stained with a fresh 5 µM dihydroethidium (DHE) staining solution (Invitrogen D1168). Slides were incubated in a dark humidified chamber at 37°C for 30-45 min to allow DHE penetration and interaction with superoxide anions. Fluorescence images were captured using a Widefield Leica AF6000 microscope equipped with the appropriate filters (excitation: 518 nm, emission: 605 nm for ethidium). Images were acquired from representative fields in each section.

### 2.9 Western blotting

Heart tissues were homogenised in RIPA buffer containing protease inhibitors (2 mM phenylmethylsulfonyl fluoride, 10 g/L aprotinin, 1 g/L leupeptin, and 1 g/L pepstatin) and phosphatase inhibitors (2 mM Na₃VO₄ and 100 mM NaF). Protein concentration was measured using a DC protein assay kit (Bio-Rad, Berkeley, CA, USA).

Nitrocellulose membranes (GE10600002 Amersham Protran, Amersham, UK) were incubated overnight at 4°C with the following primary antibodies: total OXPHOS rodent antibody cocktail (45-8099 Invitrogen; 1:5,000), TOM20 (SC-17764 Santa Cruz Biotechnology, Santa Cruz, CA, USA; 1:5,000), MMP-2 (ab37150 Abcam, Cambridge, UK; 1:1,000), and G-actin (A2066, Sigma; 1:10,000). Thereafter, membranes were rinsed and incubated with horse radish peroxidase (HRP)-conjugated secondary antibodies (W4011 (rabbit) or W021 (mouse) Promega, Madrid, Spain; 1:3,000), and protein bands were visualised using Western Blotting Luminol Reagent (Santa Cruz Biotechnology, Santa Cruz, CA, USA).

### 2.10 Images Analysis

All images were analysed using Fiji ImageJ Analysis software (ImageJ 1.54f, National Institute of Health, Bethesda, Maryland, USA).

### 2.11 Statistical analysis

Before conducting statistical analyses, data were tested for normality using the Shapiro-Wilk test. After the relevant test (parametric or not), the data were analysed to investigate potential differences due to sex. Where no significant sex-based differences were found, data were grouped based on genotype alone.

To compare genotypes, One-way ANOVA with Dunnet’s *post-hoc* test or Kruskal-Wallis test with Dunn’s post-test for multiple comparisons were used. To analyse progression over time, we used a Mixed-effects model (REML).

The statistical test applied in each case is indicated in each figure legend. Statistical details are shown in the Supplementary Tables S1-S9. Results are expressed as mean ± SEM. A *p*-value < 0.05 was considered statistically significant. All statistical analyses and graph generation were performed using GraphPad Prism 9 software.

## 3. RESULTS

### 3.1 Viability and somatic growth of CDMFS

Segregation of double heterozygous CDMFS when crossing single heterozygous MFS female mice with single heterozygous CD male mice followed the expected Mendelian ratios (χ2_(3)_=7.016, *p*=0.0714), although the number of CDMFS animals was half that of WT animals (Supplementary Table S1).

Due to significant differences in body weight between males and females, they were analysed independently. A significant effect of genotype is observed for body weight both in males (F_(3, 44)_=53.28, *p* < 0.0001) and females (F_(3, 47)_=24.67, *p* < 0.0001). Double heterozygote CDMFS mice follow a growth line highly similar to the simple heterozygote CD at two months of age (Supplementary Fig. S1). Due to somatic growth, we observed a significant effect over time for males (F_(1,847,62,17)_=106.4, *p* < 0.0001) and females (F_(2,142,74,27)_=98.19, *p* < 0.0001). However, we did not observe significant changes in the interaction between both factors (genotype and time) for males (F_(9,101)_=0.8470, *p*=0.575) or females (F_(9,104)_=1.094, *p*=0.3741), which indicates that somatic growth progresses similarly (Supplementary Fig. S1). As previously reported, CD mice had a significantly lower weight compared with WT animals ^19^, which is not the case for MFS mice ^25^. Therefore, results indicate that, for body weight, the CDMFS phenotype seems to be exclusively determined by the CD phenotype, regardless of sex. Statistical analyses are detailed in Supplementary Table S1.

### 3.2 CDMFS mice exhibit behaviour-associated alterations and brain weight like CD animals

Next, we proceeded to evaluate some cognitive outcomes, knowing that WBS patients also present certain cognitive dysfunctions, which are also mostly reproduced in the CD mouse model ^27,28^. To our knowledge, no similar behaviour studies have been previously performed in the MFS mouse model used herein. Naturally occurring behaviour was evaluated by counting the number of marbles buried in the MBT. This test was performed both in males and females at two and six months of age (Fig. 1). At two months of age, females, regardless of genotype, bury fewer marbles than males, leading to significant differences between the sexes, requiring their independent analysis (Fig. 1A). However, this statistical difference did not occur at six months of age (Fig. 1B; animals were grouped together in the analysis, but different colours distinguish males and females). Considering the nature of the MBT test, we believe that the sex difference detected in young mice is merely statistical, without any true biological significance. Double heterozygous CDMFS mice showed a significant decrease in the number of buried marbles (*p* < 0.0001, males and females at two and six months of age) (Fig. 1A and 1B). This behaviour was similar to that previously described in CD animals ^23^ and confirmed here. Single heterozygous MFS mice (males and females) performed the test similarly to WT littermates at the indicated test ages (Fig. 1A and 1B). Therefore, this specific behaviour alteration, seen only in CD mice, was transmitted to double heterozygous CDMFS mice. Statistical analyses are detailed in Supplementary Table S2.

**Figure 1.**
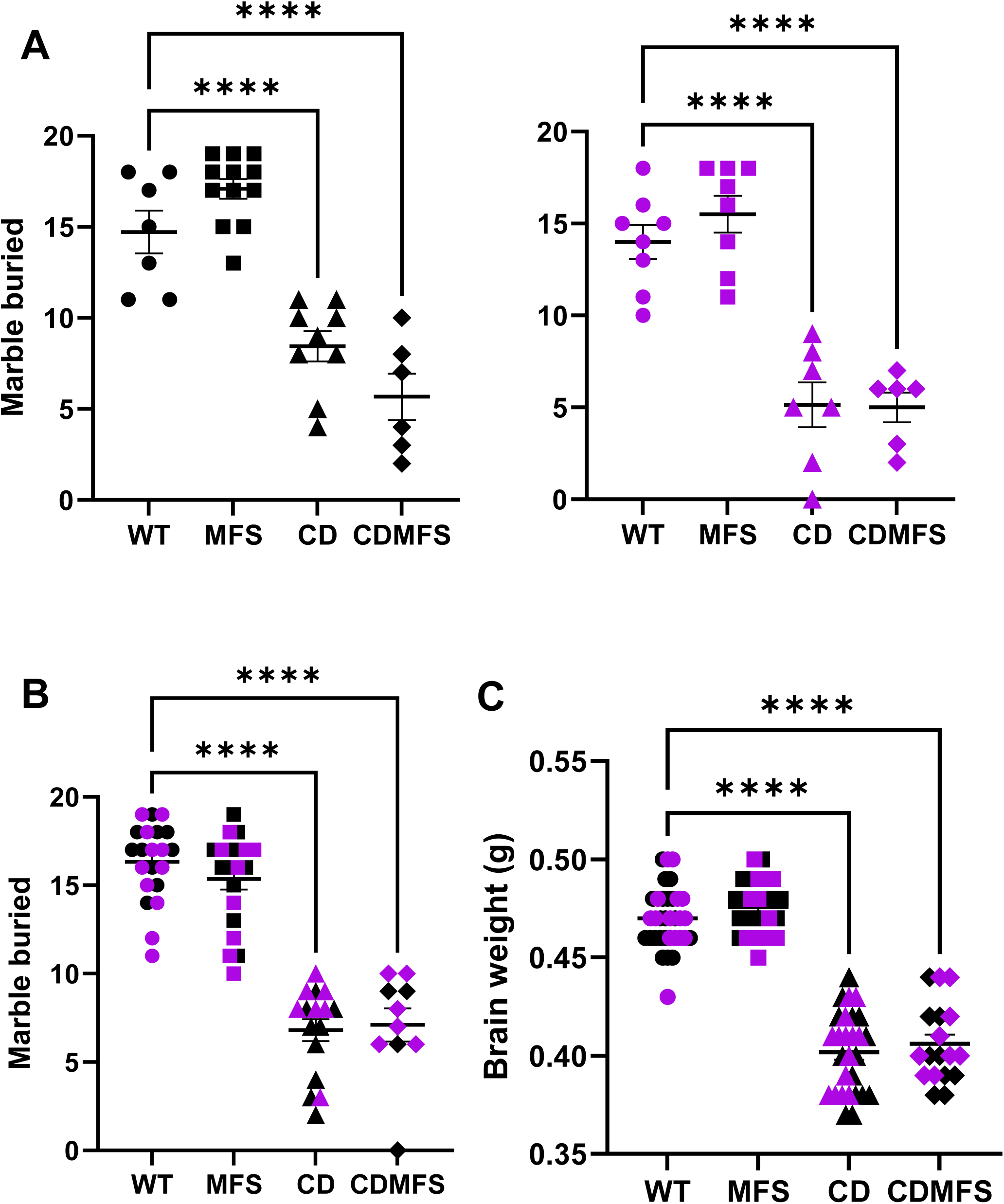
The marble burying test (MBT) and brain weight. The MBT in two (**A**) and six-month-old mice (**B**). At two months, males (black) and females (purple) were analysed separately due to the significant sex differences obtained (F_(1,55)_=5.305, *p=*0.025). At both ages, CDMFS animals hid significantly fewer marbles than their WT littermates (*p* < 0.0001) and similar to CD animals (*p* < 0.0001). MFS mice displayed behaviour indistinguishable from their WT littermates. (**C**) CDMFS animals showed significantly (*p* < 0.0001) lower brain weight compared with their WT littermates. Circles, WT; Squares, MFS; Triangles, CD; Diamonds, CDMFS. Data are expressed as the mean ± SEM. Statistical significance was calculated by a *post-hoc* Dunnett’s multiple comparisons test followed by one-way ANOVA (A) or Dunn’s multiple comparisons test followed by the Kruskal-Wallis test (B and C). \*\*\*\**p* < 0.0001

Both global and regional specific differences in brain anatomy have been reported in WBS ^32^, including reduced total brain volume, affecting white matter (18%) more than grey (11%) ^33^. We reported that brain weight was reduced in CD mice by around 9%, both in males and females ^19^. Thus, brains were extracted and weighed in all experimental groups of six-month-old mice (Fig. 1C). Results clearly showed no differences between WT and MFS animals, however, CD and CDMFS mice exhibited a severe decrease in brain weight (*p* < 0.0001). Statistical analyses are detailed in Supplementary Table S2.

Altogether, the results demonstrate no apparent contribution of *Fbn1* mutation to brain weight and MBT-associated behavioural dysfunction and, therefore, both characteristics seem to be exclusively modulated by the WBS critical region (WBSCR) deletion. In WBS individuals, deletion of *LIMK1* is associated with difficulty with visuospatial construction, part of the WBS cognitive profile. Deletions that include *GTF2I* and *GTF2IRD1* are associated with the cognitive and behavioural features of the WBS phenotype. *GTF2I* deletion is associated with intellectual disability, and *GTF2IRD1* deletion with the behavioural phenotype ^34^.

### 3.3 CDMFS vascular alterations are attributed to either elastin or fibrillin-1 haploinsufficiency or both at the same time

Next, using ultrasonography, we evaluated vascular outcomes attributable to MFS and WBS, such as their respective opposed aortopathies—aortic aneurysm in MFS and stenosis in WBS. As previously described ^35^, MFS mice showed a significantly increased AR diameter at two and six months of age (Fig. 2A and 2B, respectively). Double heterozygous CDMFS animals showed a similar AR dilatation, being indistinguishable from that of MFS mice at both ages tested (Fig. 2A and 2B). In CD mice, previous histological data showed significant aortic stenosis in three-to-four-month-old mice ^20^. We here observed a significant reduction in AR diameter, already visible in two-month-old animals (Fig. 2A). Regarding AR progression over time (Fig. 2C), statistical analysis indicates a significant effect of genotype (F_(3,125)_ =74.88, *p* < 0.0001) and time factors (F_(1,125)_ =80.24, *p* < 0.0001), but without interaction between them (F_(3,125)_ =0.6786, *p*=0.5667), indicating that the progression of AR growth is similar in all genotypes (Fig. 2C). Therefore, ultrasonography results indicate that the characteristic MFS-associated AR dilatation predominates over WBS-associated aortic stenosis. Statistical analyses are detailed in Supplementary Table S3.

**Figure 2.**
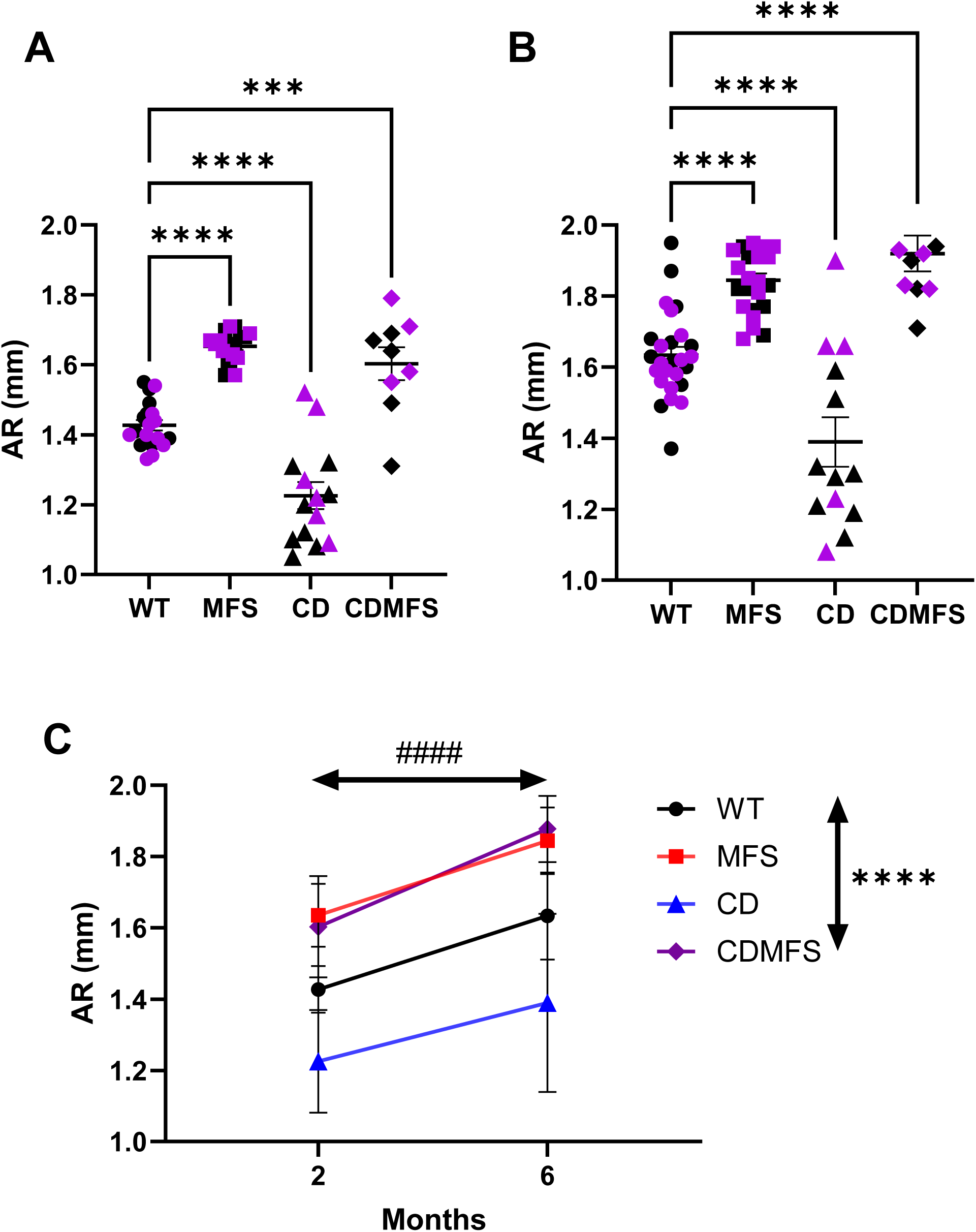
Aortic root diameter. (**A**) In two-month-old CDMFS mice, we observed a significant (*p* < 0.0001) dilatation of the aortic root (AR) similar to MFS mice. In the opposite direction, CD mice showed significant stenosis (*p* < 0.0001). These results were maintained at six months of age (**B**). (**C**) Progression of AR diameter. An increase is seen over time (F_(1,125)_=80.24, *p* < 0.0001), similar in all genotypes (interaction, F_(3,125)_=0.125, *p=*0.5667). Data are expressed as the mean ± SEM. Circles, WT; Squares, MFS; Triangles, CD; Diamonds, CDMFS; Black, males; Purple, females. Statistical significance was calculated using Dunnett’s multiple comparisons test followed by one-way ANOVA **(A and B)** and two-way Mixed effect analysis **(C)**. *Genotype effect; #time effect. \*\*\**p* < 0.001; ****, ^####^*p* < 0.0001.

Next, we analysed the histopathology of the ascending aorta in transverse sections of paraffin-embedded tissue. We measured the total area of the aorta (the media and lumen together), as well as the percentage corresponding only to the media (see Fig. 3A for representative images of each experimental group of mice). The total aortic area in CDMFS aortae was significantly reduced compared with WT littermates (*p* < 0.0001), being indistinguishable from that of CD aortae (Fig. 3B). In contrast, MFS mice exhibited a significant increase in aortic area compared with WT littermates (*p*=0.0003). Strikingly, the percentage of the media with respect to the total aortic area was significantly increased (*p* < 0.0001) in all mouse groups (Fig. 3C). In the CD model, the increase in tunica media thickness is determined by hyperplasia of VSMCs ^24^, whereas in the MFS model, a decrease in cell density has been described ^25^. Consequently, we next quantified cell density, evidencing the nuclei with DAPI staining. Strikingly, while simple heterozygous MFS and CD mice present similar results to those previously reported, CDMFS aortae showed values highly similar to these observed in WT aortae (*p*=0.655) (Fig. 3D), suggesting some type of tug-of-war between proliferation and apoptosis mechanisms. Statistical analyses are detailed in Supplementary Table S4.

**Figure 3.**
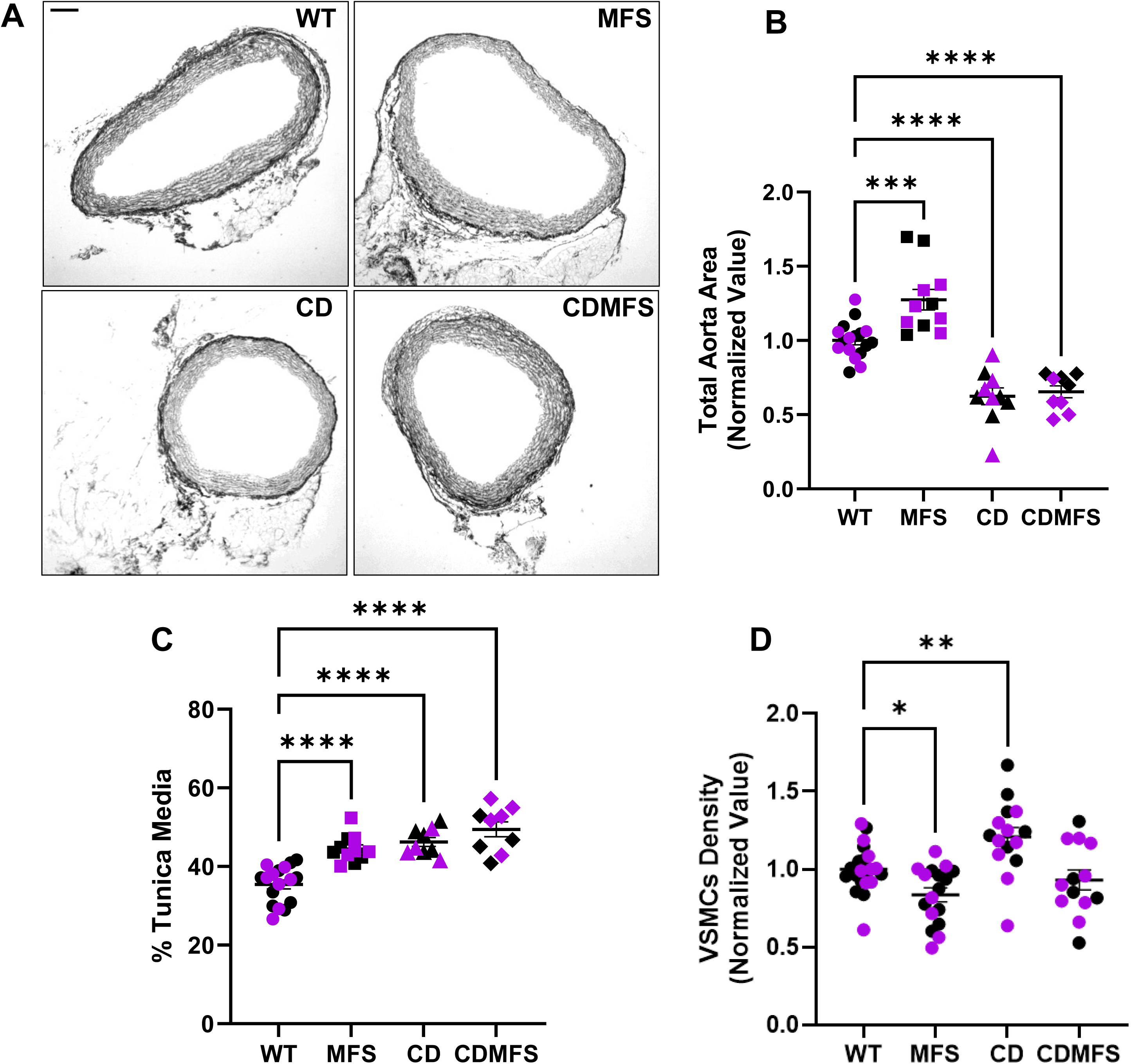
Vascular histopathology. (**A**) Representative bright-field microscopy images of paraffin-embedded ascending aorta slides. Scale bar, 100 µm. (**B**) Histogram representing the area (tunica media plus the lumen) of the ascending aorta normalised to the mean area of WT littermates. In CDMFS animals, this area was significantly smaller (*p* < 0.0001) compared with WT littermates and similar to CD. (**C**) Histogram showing the percentage of the total area occupied by the tunica media. All groups analysed showed a significant increase in this percentage. (**D**) Histogram of VSMC density (DAPI staining) relative to the mean density of WT littermates. Note that the VSMC density in CDMFS media is not significantly different (*p*=0.655) compared to WT littermates. Circles, WT; Squares, MFS; Triangles, CD; Diamonds, CDMFS; Black, males; Purple, females. Data are expressed as the mean ± SEM. Statistical significance was calculated using Dunnett’s multiple comparisons test followed by one-way ANOVA. \**p* < 0.05; \*\**p* < 0.01; \*\*\**p* < 0.001; \*\*\*\**p* < 0.0001.

To evaluate this possibility, we stained paraffin-embedded aortic sections for the presence of the characteristic proliferative marker Ki67 and apoptosis with TUNEL staining. Strikingly, results showed low activity for both processes in the media of all genotypes, with no statistical differences between them (Supplementary Fig. S2 and Supplementary Table S4).

We next examined elastic laminae integrity. Due to elastin haploinsufficiency, as expected, we observed a decrease in the autofluorescence emitted by elastic laminae of the CDMFS and CD models compared with WT littermates (Supplementary Fig. S3). Next, we evaluated elastin fibre integrity in the ascending aortae stained with Van Gieson’s method. Representative images are shown in Fig. 4A. CDMFS aortae presented more elastic fibre ruptures (*p*=0.0014) (Fig. 4B) and were more undulated (*p* < 0.0001) (Fig. 4C) compared with WT littermates. Similar results were observed in MFS aortae (*p* < 0.0001); however, this was not the case for CD aortae (*p*=0.3907), in which values for both factors (elastic breaks and undulation factor) were not statistically different from WT (Fig. 4B and 4C; the enlarged insets of the indicated aortic areas in 4A allow for improved visualisation of undulated fibres). Statistical analyses are detailed in Supplementary Table S5. The ultrastructural analysis of the aortae confirms the above results (elastic fibre amplitude and undulation level) (Supplementary Fig. S4). In addition, the characteristic continuous interaction of VSMCs with their adjacent elastic fibres can be seen to be severely altered in MFS and even more so in CDMFS aortae. VSMCs showed few and occasional interactions with neighbouring elastic fibres exhibiting a comb teeth-like phenotype. Numerous collagen fibres can be seen in the free spaces that remain between these occasional interactions (Supplementary Fig. S4). This proliferation of collagen fibres suggests a compensatory fibrotic-like response against elastic fibre disarrays.

**Figure 4.**
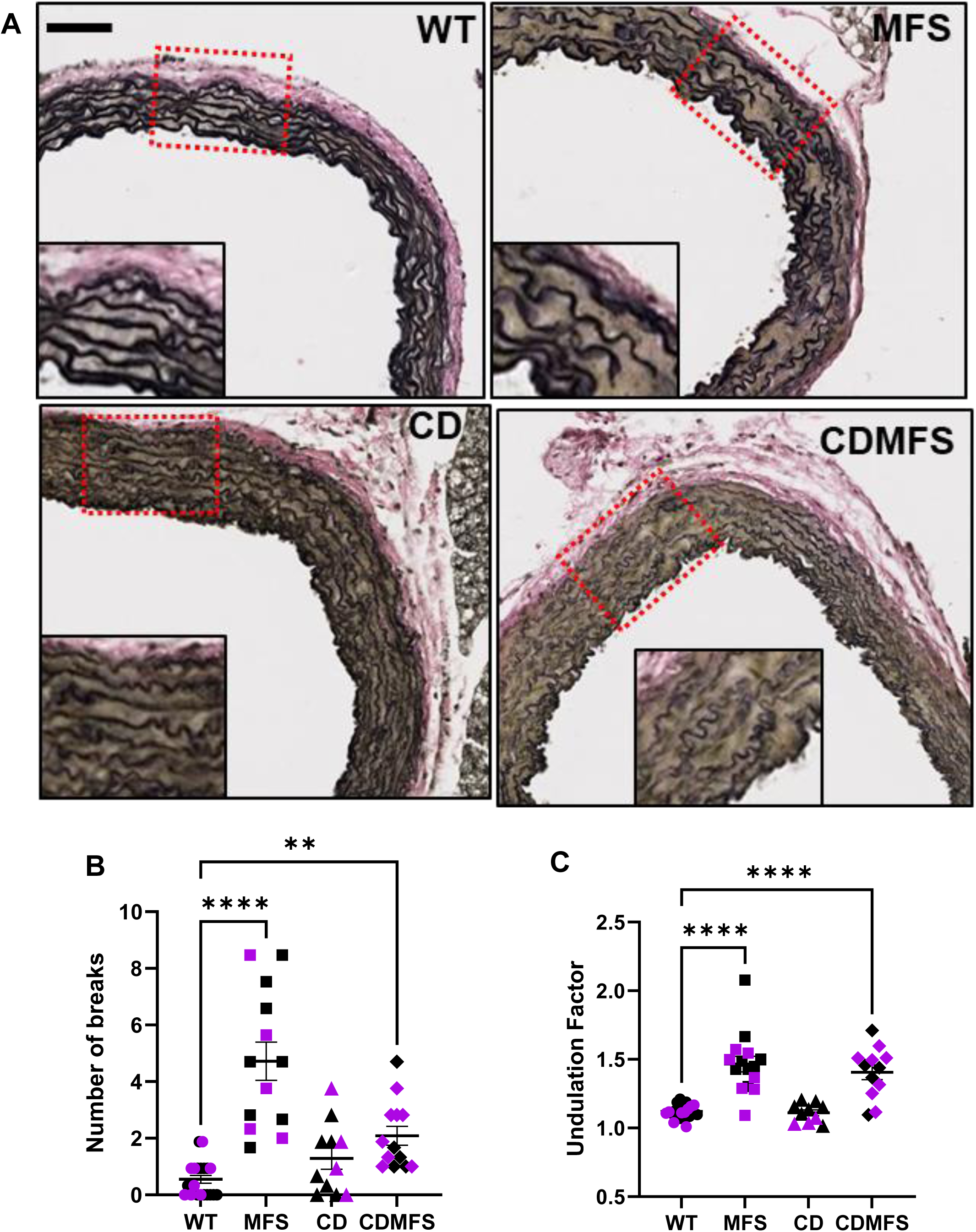
Elastic laminae integrity. (**A**) Representative pictures of ascending aorta slides stained by Van Gieson’s procedure. Scale bar, 100 µm. Insets show magnified selected areas of the indicated aortic wall (red dotted rectangles). (**B**) Histogram of large elastic fibre breaks (20 µm). CDMFS aortae showed a significant increase (*p*=0.0014) compared with WT littermates, although the mean is rather lower than in MFS tissue (4.72±0.68 *vs*. 2.09±0.33). (**C**) Assessment of the elastic laminae undulation factor. CDMFS elastic fibres showed a significant increase in the undulation factor (*p* < 0.0001), similar to MFS. Circles, WT; Squares, MFS; Triangles, CD; Diamonds, CDMFS; Black, males; Purple, females. Data are expressed as the mean ± SEM. Statistical significance was calculated by Dunn’s multiple comparisons test followed by the Kruskal Wallis test (**B**) and Dunnett’s multiple comparisons test followed by one-way ANOVA (**C**). \*\*\**p* < 0.001; \*\*\*\**p* < 0.0001.

In sum, these results indicate that, with the exception of the reduced thickness of the elastic fibres (as a result of the elastin haploinsufficiency), the CDMFS aortic media disarrays are highly similar to those of the MFS phenotype, indicating little contribution from the CD aortic phenotype.

### 3.4 The higher blood pressure observed in CDMFS mice is attributable to the CD phenotype

It is well-known that, unlike MFS patients and mice, WBS patients and CD mice have elevated SBP ^19,36–38^. Therefore, we evaluated this cardiovascular outcome in CDMFS animals over time (two and six months of age) (Fig. 5). CDMFS mice already present increased SBP at two months of age (*p*=0.0014) (Fig. 5A); at six months, this difference is even more pronounced (*p* < 0.0001) (Fig. 5B). In the progression of SBP (Fig 5C), a time effect is observed (F_(1,61)_=67.24, *p* < 0.0001) without a genotype-time interaction (F_(3,61)_=0.2325, *p*=0.8734) because the increase occurs equally across all genotypes with age (Fig. 5C). Therefore, the characteristic elevated SBP observed in CD mice also occurs in double heterozygous CDMFS animals with no apparent modulation by the MFS phenotype. Statistical analyses are detailed in Supplementary Table S6.

**Figure 5.**
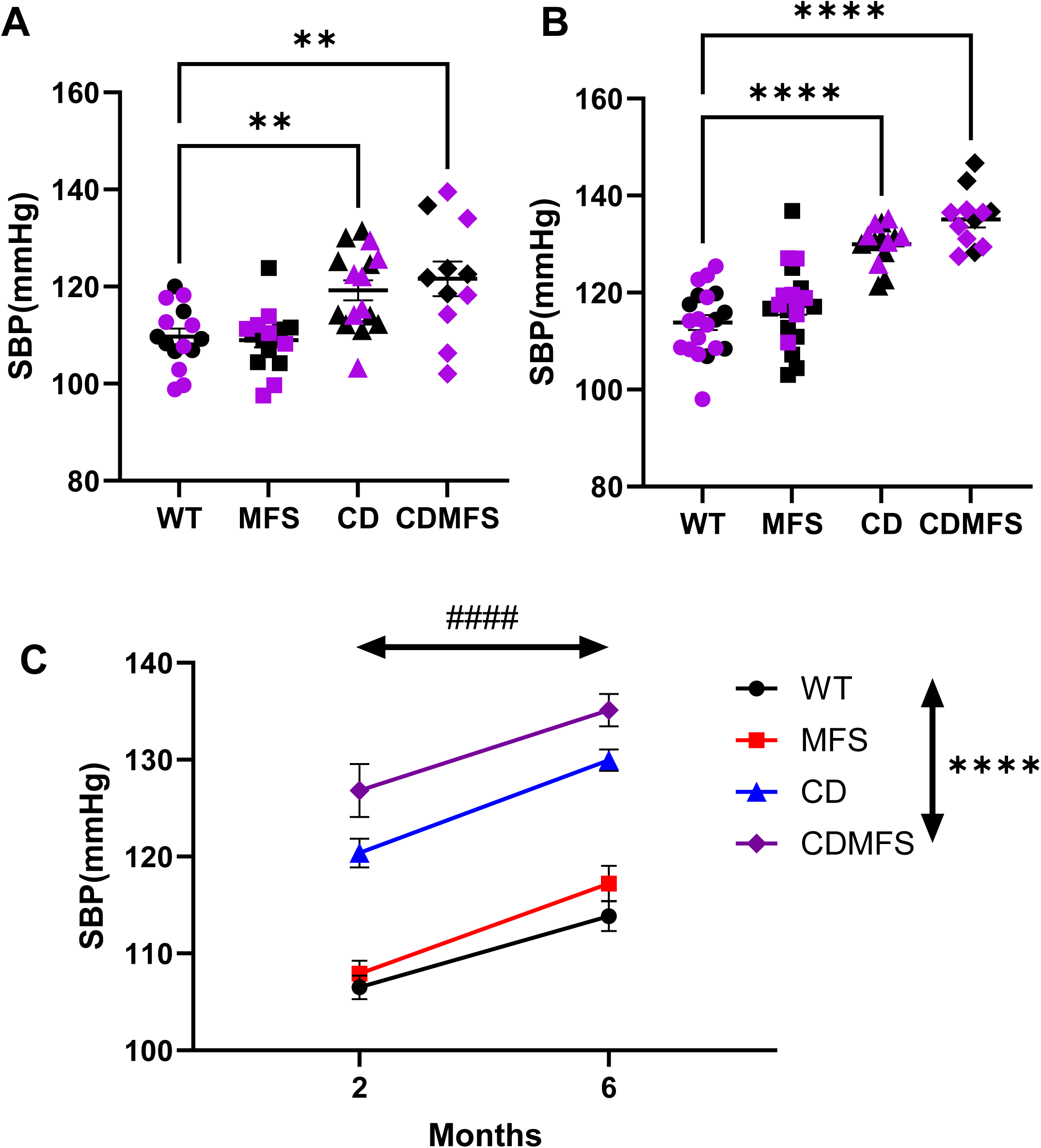
Systolic blood pressure analysis. SBP was measured in two- (**A**) and six-month-old mice (**B**) in the conscious state using the indirect tail-cuff method. CDMFS mice showed significantly increased SBP compared with WT littermates (*p*=0.0014 (A) and *p* < 0.0001 (B)**. (C)** SBP progression over time. A significant effect of genotype and time was observed without any significant interaction between factors (F_(3,61)_=0.2325, *p*=0.8734). Circles, WT; Squares, MFS; Triangles, CD; Diamonds, CDMFS; Black, males; Purple, females. Data are expressed as the mean ± SEM. Statistical significance was calculated by a *post-hoc* Dunnett’s multiple comparisons test followed by one-way ANOVA (**A**, **B**), and two-way Mixed effect analysis (**C**). *Genotype effect; #time effect. \*\**p* < 0.01; ****, ^####^*p* < 0.0001.

### 3.5 Cardiac dysfunction and hypertrophy occur in MFS, CD, and CDMFS mice

In parallel to the aorta ultrasonography, we analysed a variety of cardiac parameters clearly associated with MFS or WBS phenotypes. In CD animals, cardiac dysfunction is associated with cardiac hypertrophy ^21,39^. Therefore, we examined the heart/body weight ratio in all experimental groups at six months of age (Fig. 6A). All mutant genotypes had significantly increased ratios compared with WT littermates, which is indicative of damaged heart functionality and cardiac hypertrophy both in males and females. This is also supported by the increased thickness at end-diastole of the IVSd and the LVWPd in six-month-old mice (Supplementary Fig. S5 and S6; Statistical analyses are detailed in Supplementary Table S7).

**Figure 6.**
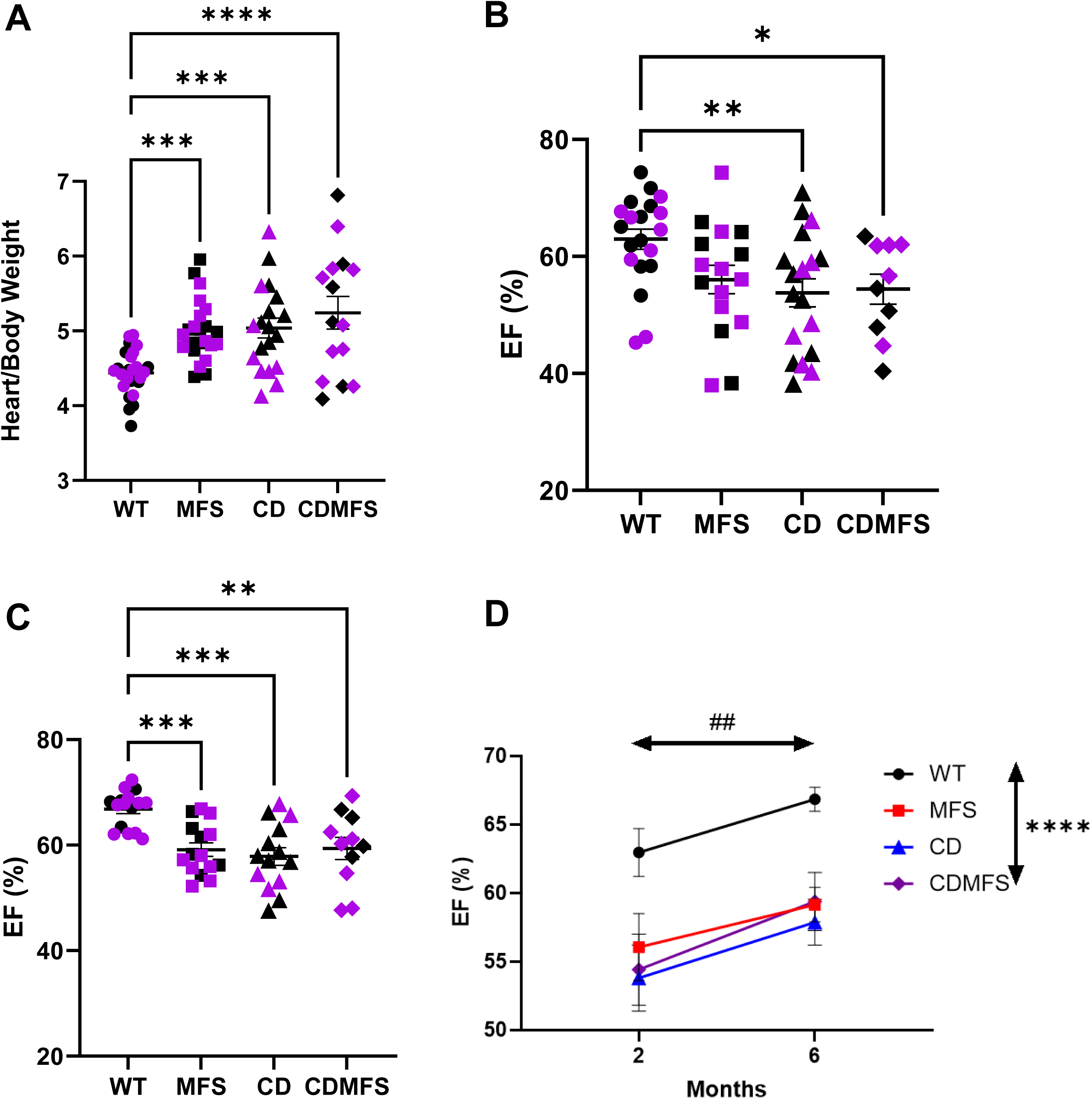
Cardiac pathology**. (A)** Significantly increased heart/body weight ratios in all genotypes. (**B and C**) Ejection fraction (EF) (%) in two (B) and six-month-old mice (C). All genotypes revealed a significant decrease in the EF. MFS animals do not reach significant values (*p*=0.073) at two months of age (B); however, the dysfunction is clearly established (*p*=0.0007) at six months (C). (**D**) EF progression over time. No interaction between genotype and time was observed (F_(3,113)=_0.721, *p*=0.5414). Circles, WT; Squares, MFS; Triangles, CD; Diamonds, CDMFS; Black, males; Purple, females. Data are expressed as the mean ± SEM. Statistical significance was calculated by a *post-hoc* Dunnett’s multiple comparisons test followed by one-way ANOVA (A, B, C) and two-way Mixed effect analysis. *Genotype effect; # time effect) (D). \**p* < 0.05; **, ^##^ *p* < 0.01; \*\*\**p* < 0.001; \*\*\*\**p* < 0.0001.

As a functional indicator of cardiac function, we next evaluated the ejection fraction (EF) in all experimental mouse groups at two (Fig. 6B) and six months of age (Fig. 6C). In all mutant groups, we observed a reduced EF compared with WT littermates, whereas in two-month-old MFS mice, the reduction was not yet statistically significant (*p*=0.073) (Fig. 6A). In the EF progression analysis (Fig. 6D), we observed a significant effect of genotype (F_(3,113)_=17.64, *p* < 0.0001), in agreement with the data analysed at different times (Fig. 6A and 6B). There was also a significant effect of time (F_(1,113)_=12.4, *p*=0.0006), with increased values at six months of age (Fig. 6D), probably because of the increased body weight since there is no interaction between the two factors (genotype/time; F_(3,113)_=0.7211, *p*=0.5414). Therefore, double heterozygous CDMFS mice exhibit a heart failure-like phenotype revealed by a reduced EF, which was also observed in single heterozygous MFS and CD mice (statistical analyses are detailed in Supplementary Table S7). The significant increase observed in EF over time (Fig. 6C) might be attributable to intrinsic physiological changes associated with age.

Finally, we evaluated fibrosis in the cardiac tissue after staining with Sirius red. No differences were observed between the four genotypes (Supplementary Fig. S7). Statistical analyses are detailed in Supplementary Table S7

### 3.6. CDMFS shows mitochondrial cardiomyocyte dysfunction and redox stress

Redox stress participates to a variable but significant extent in both MFS and WBS cardiovascular injuries ^40–45^. Dysfunctional mitochondria are essential in redox stress generation and have been previously reported in both disorders ^46,47^. Thus, we next evaluated mitochondrial functionality by analysing the protein expression levels of OXPHOS complexes. Western blot experiments showed that MFS, CD, and CDMFS cardiomyocytes had significantly reduced protein levels of all complexes (F_(3, 27)_=26.15, *p*<0.0001, effect of genotype) (Fig. 7A). This result suggests mitochondrial dysfunction in cardiomyocytes of both diseases. Ultrastructural evaluation of cardiac tissue also showed morphological alterations in their characteristic rounded mitochondria located between cardiac fibres. CD mitochondria were aligned and joined together at a single point (Supplementary Fig. S8). This peculiar morphology was also seen in CDMFS and, to a lesser extent, MFS cardiomyocytes, though not as pronounced as in CD cells. This result suggests, in agreement with previous CD mice data, a potential shift towards mitochondrial fission, as previously indicated ^47^.

**Figure 7.**
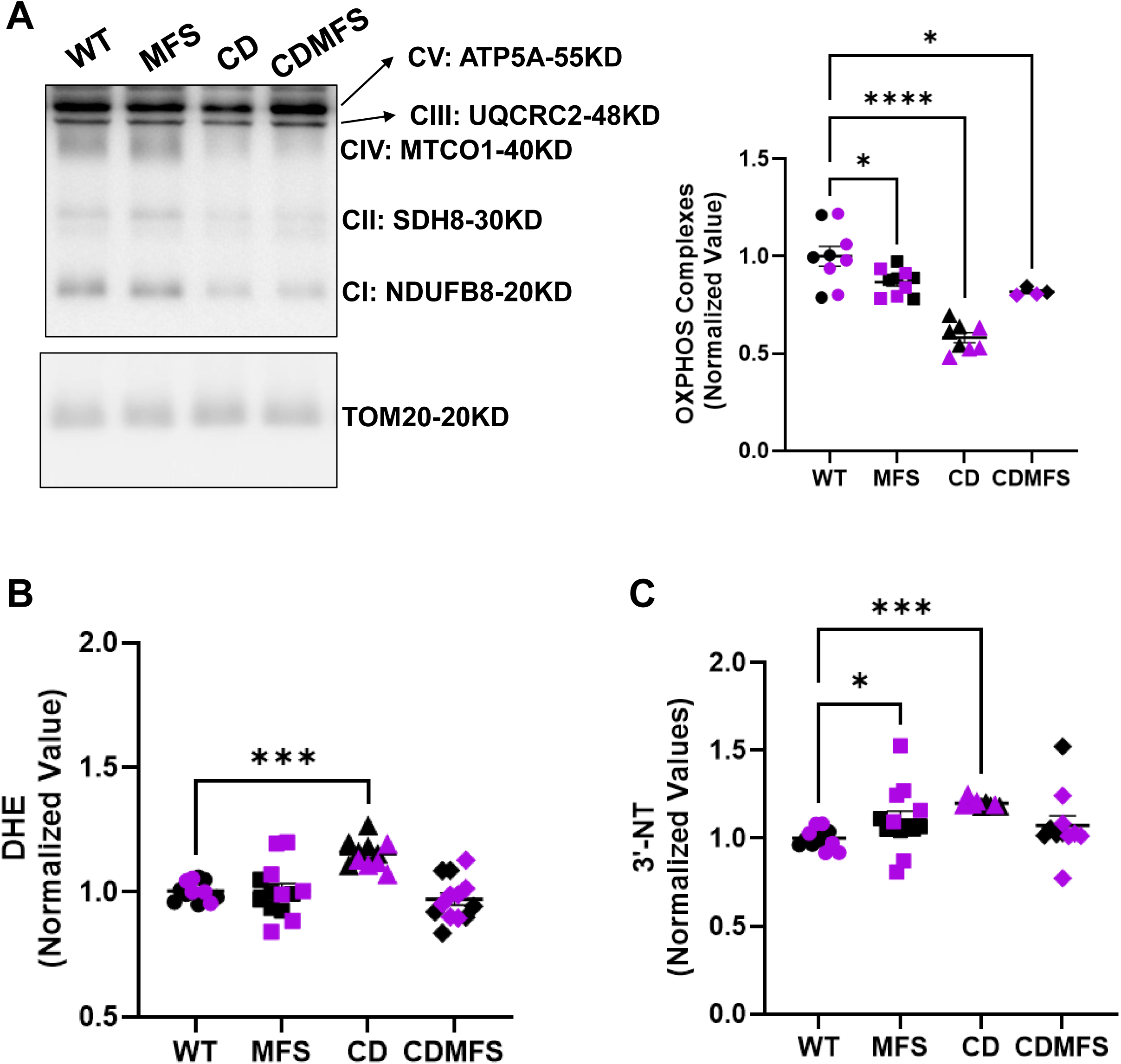
Redox stress analysis. (**A**) On the left, a representative western blot image for OXPHOS protein complexes revealed in cardiomyocytes with a specific COKTAIL antibody from the four experimental groups; on the right, a histogram showing the quantitative analysis of OXPHOS complex proteins normalised against mitochondrial protein TOM20. A significant reduction in these complexes is observed in all mutant genotypes, although it was more exacerbated in CD animals. The oxidative stress markers DHE (**B**) and 3-nitrotyrosine residues (3-NT) (**C**) were quantified by fluorescence in the left ventricles. No significant differences were observed in CDMFS animals compared to WT samples. *p*=0.6024 in (B) and *p*=0.5956 in (**C**). Circles, WT; Squares, MFS; Triangles, CD; Diamonds, CDMFS; Black, males; Purple, females. Data are expressed as the mean ± SEM. Statistical significance was calculated using Dunnett’s multiple comparisons test followed by one-way ANOVA. (**A and B**) and Dunn’s multiple comparisons test followed the Kruskal-Wallis test (**C**). \**p* < 0.05; \*\*\**p* < 0.001; \*\*\*\**p* < 0.0001.

Thereafter, we evaluated redox stress markers by DHE (Fig. 7B) and 3’-NT (Fig. 7C) fluorescent staining of paraffin-embedded sections of the left ventricle of six-month-old mice from the experimental groups. Double heterozygous CDMFS mice showed no differences with respect to their WT siblings for any of the markers analysed (*p*=9954 for DHE and *p*=0.6778 for 3’-NT). However, in agreement with published results ^21,24^, we observed a significant increase in both markers in CD mice (*p*=0.0001 for DHE and 3’-NT), while MFS only showed increased values for 3’-NT residues (*p*=0.0344). This result suggests some protective cardiomyocyte anti-redox stress response in MFS mice that is not produced by elastin deficiency. Statistical analyses are detailed in Supplementary Table S8.

### 3.7 Unlike CD, both MFS and CDMFS hearts show increased protein expression levels of MMP-2

Changes in metalloproteinase 2 (MMP2) protein expression have been reported as a potential biomarker of cardiac damage in myocardial infarction and hypertension-associated heart disease ^48^. In addition, the MFS mouse aorta shows increased MMP-2 levels ^49^, however, it is unknown if this also occurs in the heart. Thus, we examined MMP2 protein levels in the hearts of all experimental mouse groups at six months of age (Fig. 8). The MMP2/actin ratio significantly increased in CDMFS (*p*=0.0312) and MFS (*p*=0.0006) cardiac samples but not in CD (*p*=0.4095). This result indicates that MMP2 overexpression in CDMFS cardiomyocytes is primarily determined by fibrillin-1 mutation and not by elastin haploinsufficiency. Statistical analyses are detailed in Supplementary Table S9.

**Figure 8.**
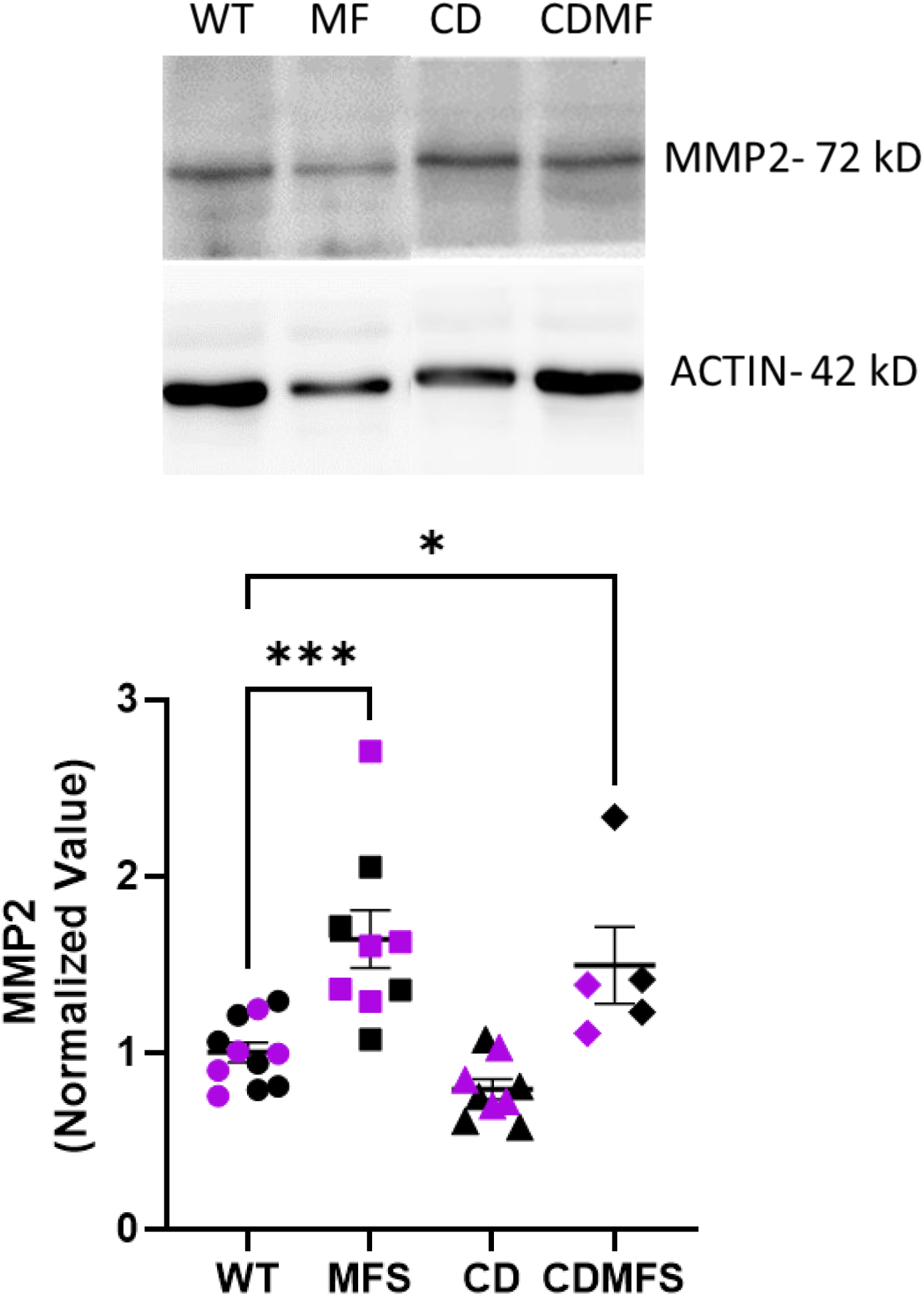
MMP-2 levels. Representative western blot image for MMP-2 protein expression levels. Below is a quantitative analysis of MMP-2 normalised to G-Actin and subsequently expressed relative to the mean of WT littermates. Note the significant increase (*p*=0.0312) in MMP-2 protein levels both in MFS and CDMFS left ventricles. CD samples showed no change in MMP-2 protein levels compared with MFS and CDMFS hearts. Circles, WT; Squares, MFS; Triangles, CD; Diamonds, CDMFS; Black, males; Purple, females. Data are expressed as the mean ± SEM. Statistical significance was calculated using Dunnett’s multiple comparisons test followed by one-way ANOVA. \**p* < 0.05; \*\*\**p* < 0.001.

## 4. DISCUSSION

Marfan syndrome (MFS) and Williams-Beuren (WBS) syndrome are two rare genetic diseases that primarily affect extracellular matrix organisation and dynamics due to the appearance of genetic variants of the fibrillin-1 glycoprotein in the case of MFS and elastin protein haploinsufficiency from chromosomic deletion in WBS. It is curious that despite both being molecular components essential for elastic fibre assembly, their genetic-based alterations generate opposed cardiovascular phenotypes, particularly at the aortic level, i.e., aortic aneurysm (MFS) and stenosis (WBS). In this study, we aimed to discern in detail the impact of elastin haploinsufficiency and fibrillin-1 mutation in the development of the cardiovascular phenotype in a double heterozygous model of MFS (*Fbn1^C1041G/+^*) and WBS (CD), hereafter CDMFS. We aimed to determine the extent to which the resulting cardiovascular phenotype in this mixed model is interdependent on both. The results reported provide interesting insights into the impact and prevalence of each elastic molecular component over the other, depending on the cardiovascular outcome studied. Thus, the main results observed in CDMFS mice are detailed as follows:

i. The elastin haploinsufficiency-based phenotype (CD model of WBS) is prevalent over that of MFS regarding brain-tested dysfunction as indicated by the MBT, which could probably be linked to reduced brain and body weights. This is not surprising as it was previously reported in WBS patients ^33,50^ and CD mouse models ^19,33,49^. It is thought to be a consequence of the additive effects of several genes in the deleted region, both proximal (PD) and distal (DD, including *ELN*) ^51^. The PD region is responsible for the anxiety-related response, whereas the DD region may also contribute to the anxiety response, at least in males ^51^. Therefore, characteristic altered behavioural responses seen only in CD mice were transmitted to double heterozygous CDMFS animals. The contribution of the additional fibrillin-1 mutation, in this case, seems to be null. Obviously, we cannot dismiss the fact that other genetic variants of fibrillin-1 and/or another MFS mouse model could modulate or exacerbate this or another behaviour. This certainly warrants further study, particularly more so knowing that MFS patients also present psychological dysfunction ^52^.
ii. Regarding the cardiovascular outcomes examined, results in CDMFS mice are variable concerning the contributions of the parental WBS and MFS phenotypes, which range from exacerbated to cancelling each other or at least mitigating the impact of the other. When analysing the AR by echocardiography, the CDMFS aorta showed the characteristic aneurysm of MFS, the results being indistinguishable from that obtained in MFS mice. This is just the opposite of what occurs in CD mouse and WBS patient aortopathy (aortic stenosis) (compare Fig. 2A and B). Strikingly, in the histopathological examination of the integrity of aortic elastic fibres, CDMFS mice presented the opposite result to MFS aorta and, therefore, a highly similar phenotype to CD aorta, with an aortic wall with much more continuous elastic fibres almost devoid of elastic breaks, being closer to the WT aortic phenotype (Fig. 2A and B). Concomitantly to these observations, MFS and CDMFS elastic fibres are more undulated than WT and CD ones, which correlates with non-pressurised aortae of the histological preparations (smaller diameter) compared with the dilatation presented in the ultrasound tests (intermediate pressure aortas under anaesthesia). These results also agree with previously reported mechanical studies in which elastin and fibrillin-1 were postulated to have distinct roles in vessel distensibility^29^. From this study, the authors indicate that elastin provides elastic recoil (heterozygous *Eln^+/−^* mice aorta have a smaller diameter at 0 mmHg; evaluated by histology), and fibrillin-1 provides tensile strength (heterozygous *Fbn1^+/−^* mice aorta have a significantly larger diameter at 75-100 mmHg pressure; evaluated by ultrasound). Another consequence of these results is that the aortic dilatation (visualised by ultrasonography) does not necessarily need to be accompanied by elastic fibre ruptures. It is possible that the normalisation of the aortic wall in terms of VSMC density and, in turn, the media area seen in CDMFS and CD mice, to some extent, preserves the morphological integrity of elastic fibres. Ultrastructural analysis of the aortic media of the aorta from the experimental groups shows that the CDMFS aorta has thinner elastic laminae compared with MFS aortae. In any case, the structural organisation of the media in MFS, CD, and CDMFS is invariably affected, in which VSMCs clearly show altered morphology and a high degree of disconnection with adjacent fibres and collagen proliferation between cells and elastic fibres, which, in the case of MFS, explains the known elevated aortic stiffness ^53–55^.

All mutant models show thickening of the aortic tunica media compared with their WT counterparts. The cause of this thickening differs between models. On the one hand, in the MFS aorta, it is caused by the increased presence of glycans and collagens that accompanies the reduction in VSMC ^56–58^. Conversely, the main cause of increased media thickness in CD aortae is VSMC hyperplasia. Strikingly, evaluation of the proliferation and apoptosis levels in aortae of the four genotypes revealed them to be very low and balanced.

The characteristic elevated SBP observed in WBS patients ^37^ and CD mice [19] has always been attributed to SVAS due to elastin deficiency (*Eln^+/−^* mice are also hypertensive) ^29,59^. We observed that double heterozygous CDMFS animals are also hypertensive. In line with previous data ^29^, our results support the idea that elastin plays a greater role than fibrillin-1 in establishing baseline blood pressure. In the same study, authors associated hypertension with a significant increase in left ventricular (LV) wet weight in double heterozygous *Eln^+/-^-Fbn1^+-/^; ELN/FBN)*, and with a remarkable trend towards the same increase in *Eln^+/−^*mice ^29^. In our study, note that at a later age (six months) and despite MFS mice having normal SBP, fibrillin-1 mutations contributed to the exacerbated SBP seen in CDMFS mice compared with CD animals (Fig. 5B). It is possible that the apparently “healthy” SBP of MFS mice (and perhaps in patients as well) could be considered “unhealthy” as the values obtained seem to predispose the animals to the altered cardiac parameters associated with hypertension, such as the reported thickening of the intraventricular septum ^60^. We also observed a significant increase in the wet weight of the heart in CDMFS mutants, like that presented by each of the simple heterozygous MFS and CD mice in accordance with previous observations in WBS and MFS patients ^16,61^. This result could be associated with the diminished EFs observed in all mutant mice groups. Note that cardiac injuries were not particularly aggravated in CDMFS mice compared with CD and MFS independently, suggesting that there is no additive or synergistic damage between both elastin and fibrillin-1 haploinsufficiency. Since oxidative stress contributes to cardiovascular damage in rare genetic aortic diseases ^44^, we evaluated the functional level of mitochondria as the main source of reactive oxygen species. All mutant genotypes presented deficiencies, to different extents, in the protein levels of respiratory complexes (OXPHOS) compared with their WT counterparts. In the case of CD and MFS animals, our results confirm previously reported observations ^46,47^, however, the damage in CD cardiomyocytes is more severe than in MFS cells. Strikingly, CDMFS cardiomyocytes show similar levels to MFS cells. Since mitochondria are of maternal inheritance, this milder phenotype can be explained as mitochondria were derived from the MFS mother. However, when we analyse redox stress markers (DHE and 3-NT), we observe that only CD mutant hearts present significantly elevated values, while MFS and CDMFS animals present values similar to WT. CDMFS results suggest some protective cardiac anti-redox stress effect triggered by fibrillin-1 over elastin haploinsufficiency. This could be related to a higher endogenous antioxidant machinery response (e.g., NRF2) that is seriously damaged in CD cardiomyocytes ^21^.

One indirect source of mitochondrial dysfunction could be MMP2 activation ^62^. Our results show significantly increased MMP2 levels in the cardiac tissue of both MFS and CDMFS models. MMP2 is a TGF-β-induced metalloproteinase localised to cardiomyocytes ^63^. It is known that full-length MMP2 is quickly activated during oxidative stress ^64^ and elevated MMP2 protein and activity levels have been described in the ascending aorta of the MFS model ^65^; these levels are reduced after treatment with losartan ^66^ and the antioxidant allopurinol ^49^. Taken together, our data imply that the cardiac dysfunction observed in CDMFS would be more attributable to fibrillin-1 than elastin haploinsufficiency.

We are aware that our study has its limitations. It would be of interest to study other outcomes characteristic of each disease, such as skeletal, ocular, and cognitive injuries, but we have focused on the most critical ones for the survival and quality of life of patients. In contrast to CD, the MFS mouse model is only representative of one type of mutation among the many that occur in the human gene. We, therefore, cannot discard that other phenotypic characteristics and studied outcomes could behave differently in the presence of the CD-associated deletion. Finally, for comparative purposes with the study carried out with the *Eln^+/-^-Fbn1^+/-^*mouse model [29], it would be of interest to evaluate the reactivity of the CDMFS aorta by myography.

In summary, the results of a mixed transgenic mouse between MFS and WBS phenotypes provide interesting new insights into the interdependence of fibrillin-1 and elastin proteins in cardiovascular and brain outcomes. Brain dysfunctions seem to be exclusively attributed to the chromosomic microdeletion of WBS and are probably not associated with elastin haploinsufficiency. Cardiovascular disorders (in the aorta and heart) show a high level of independence with respect to the aortic dilatation but complementarity concerning the organisation of the aortic wall and its disarray. At the cardiac level, fibrillin-1 haploinsufficiency seems to have a greater impact than elastin. Finally, BP and cardiopathy results in CDMFS mice indicate that both are not necessarily linked to each other as is usually thought, and fibrillin-1 haploinsufficiency could participate, somehow, in enhancing the characteristic intrinsic hypertension of the CD model. Therefore, the distinct phenotypes and the lack of exacerbation of the single phenotypes in the CDMFS mouse show that both syndromes encompass more than an elastin fibre defect, which should divert the focus to other contributors of these diseases. Supplementary Table S10 shows a summary of the main results obtained in our study.

## Author Contributions

Substantial contributions to the conception/ design of the work: VC-GE

The fine-tuning, acquisition, analysis, drafting, and data interpretation of the work: IRR-LS-JRC-AA-VC-GE

Revising the work critically for important intellectual content: IRR-APD GE -VC-Final approval of the version to be published: IRR-LS-JRC-AA-VC-GE

Agreement to be accountable for all aspects of the work in ensuring that questions related to the accuracy or integrity of any part of the work are appropriately investigated and resolved: IRR-LS-JRC-AA-VC-GE.

For the preparation and writing of this article, the authors declare that no IA application has been used.

## Acknowledgements

We thank Maria Encarnación Palomo and the Advanced Microscopy Unit of the University of Barcelona for technical assistance, and Helena Kruyer for editorial work. G.E. is particularly grateful to SIMA (Spanish Association of Patients with Marfan Syndrome and Related Diseases), MARFANS (l’Association Française des Syndromes de Marfan, Loeys-Dietz et Apparentés) and patients for their economic contributions to the research in this lab.

## SUPPLEMENTARY TABLES

**Table S1.** Statistical Analysis of Viability and Body Weight. Detailed statistical analysis of data presented in Section 3.1 and shown in Supplementary Figure S1

**Table S2.** Statistical Analysis of the Marble Burying Test and Brain Weight. Detailed statistical analysis of data discussed in Section 3.2 and shown in Figure 1

**Table S3.** Statistical Analysis of Aortic Root Diameter. Detailed statistical analysis of data discussed in Section 3.3 and shown in Figure 2

**Table S4.** Statistical Analysis of Vascular Histology. Detailed statistical analysis of data discussed in Section 3.3 and shown in Figure 3 and in Supplementary Figure S2.

**Table S5.** Statistical Analysis of Elastic Laminae Integrity. Detailed statistical analysis of data discussed in Section 3.3 and shown in Figure 4 and Supplementary Figure S3.

**Table S6.** Statistical Analysis of Systolic Blood Pressure. Detailed statistical analysis of data discussed in Section 3.4 and shown in Figure 5.

**Table S7**. Statistical Analysis of Cardiac Dysfunction. Detailed statistical analysis of data discussed in Section 3.5 and shown in Figure 6 and Supplementary Figures S5, S6, and S7.

**Table S8**. Statistical Analysis of Redox Stress Markers. Detailed statistical analysis of data discussed in Section 3.6 and shown in Figure 7.

**Table S9.** Statistical Analysis of MMP-2 Expression. Detailed statistical analysis of data discussed in Section 3.7 and shown in Figure 8.

**Table S10.** Summary of the analysis of the altered outcomes observed in CDMFS mice.

---

**Figure S1.**
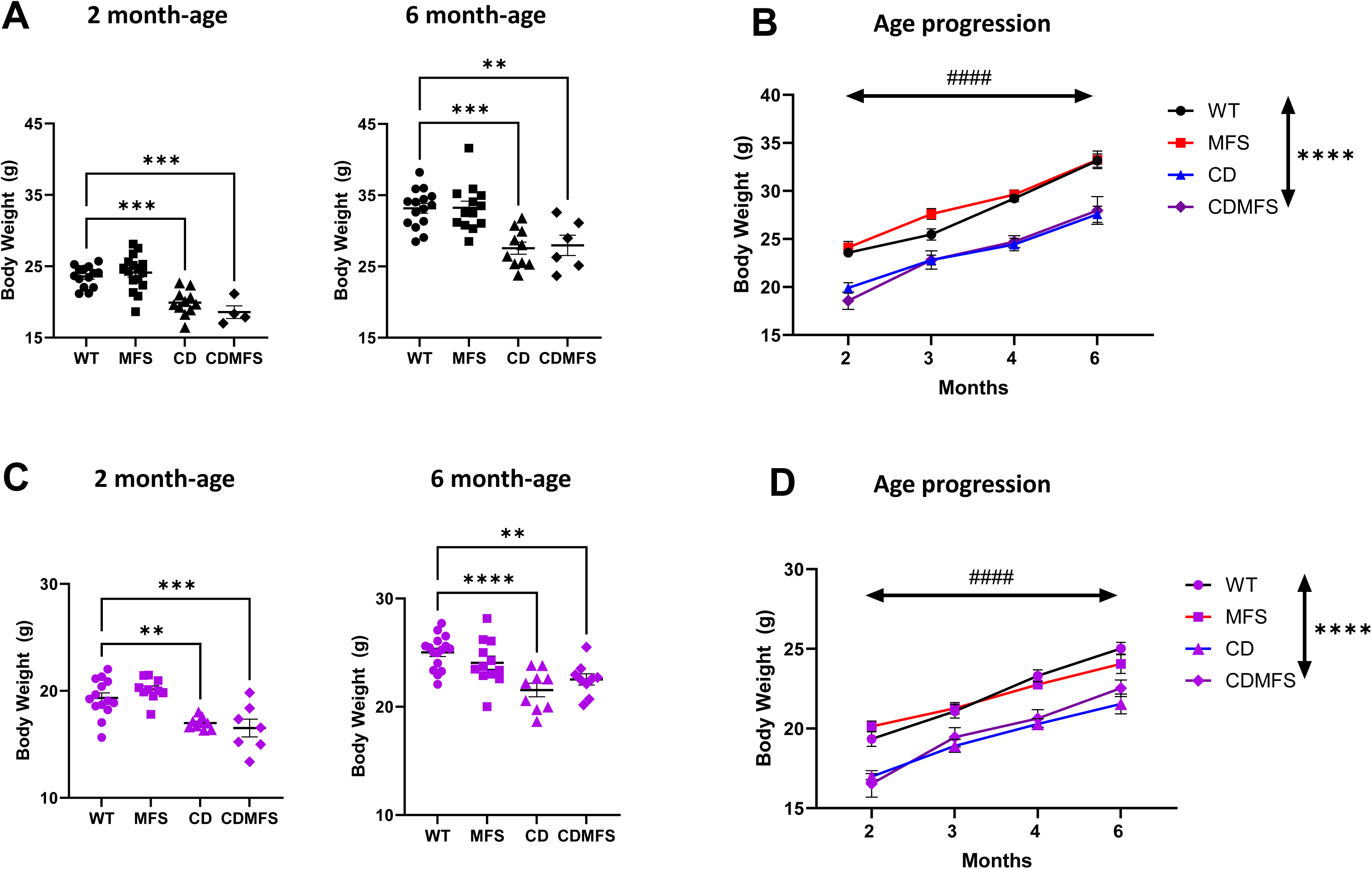
Body weight. All animals were weighed at two and six months of age **(A, C)**. The weight of the animals was analysed by sex due to significant differences in this factor. CDMFS animals (males and females) showed a significantly lower weight at two months of age, which was maintained until six months. No significant effect interaction between genotype and time was observed for males **(B, D)**, showing the progression of weight over time in males (F_(9,101)=_0.847, *p=*0.575) and females **(D)** (F_(9,104)=_1.094, *p=*0.374). Circles, WT; Squares, MFS; Triangles, CD; Diamonds, CDMFS. Data are expressed as the mean ± SEM. Statistical significance was calculated by Dunnet’s comparison test followed by an ordinary one-way ANOVA (**A, C)** and two-way Mixed effect analysis **(B, D)**. *Genotype effect; # time effect. \*\**p* < 0.01, \*\*\**p* < 0.001, ****, ^####^*p* < 0.0001.

**Figure S2.**
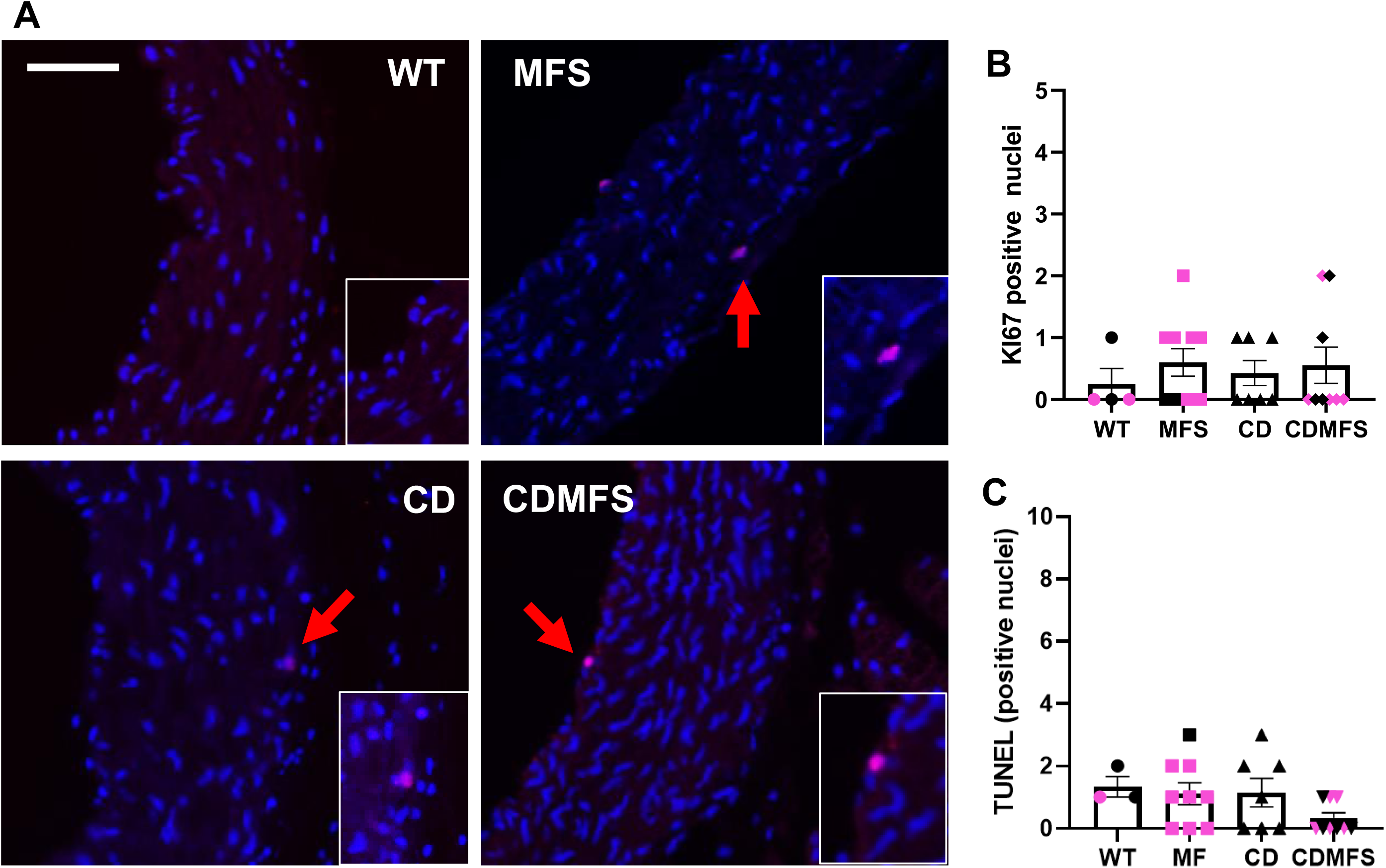
Proliferation and apoptosis in the aortic media. (**A and B**) Ki67 immunofluorescence **(A)** and quantitative analysis **(B)** in WT, MFS, CD, and CDMFS aortae. Red arrows indicate co-staining for Ki67 and DAPI, whose images are enlarged in the respective insets. Quantitative analysis shows no differences between genotypes. Circles, WT; Squares, MFS; Triangles, CD; Diamond, CDMFS. Data are expressed as the mean ± SEM.

**Figure S3.**
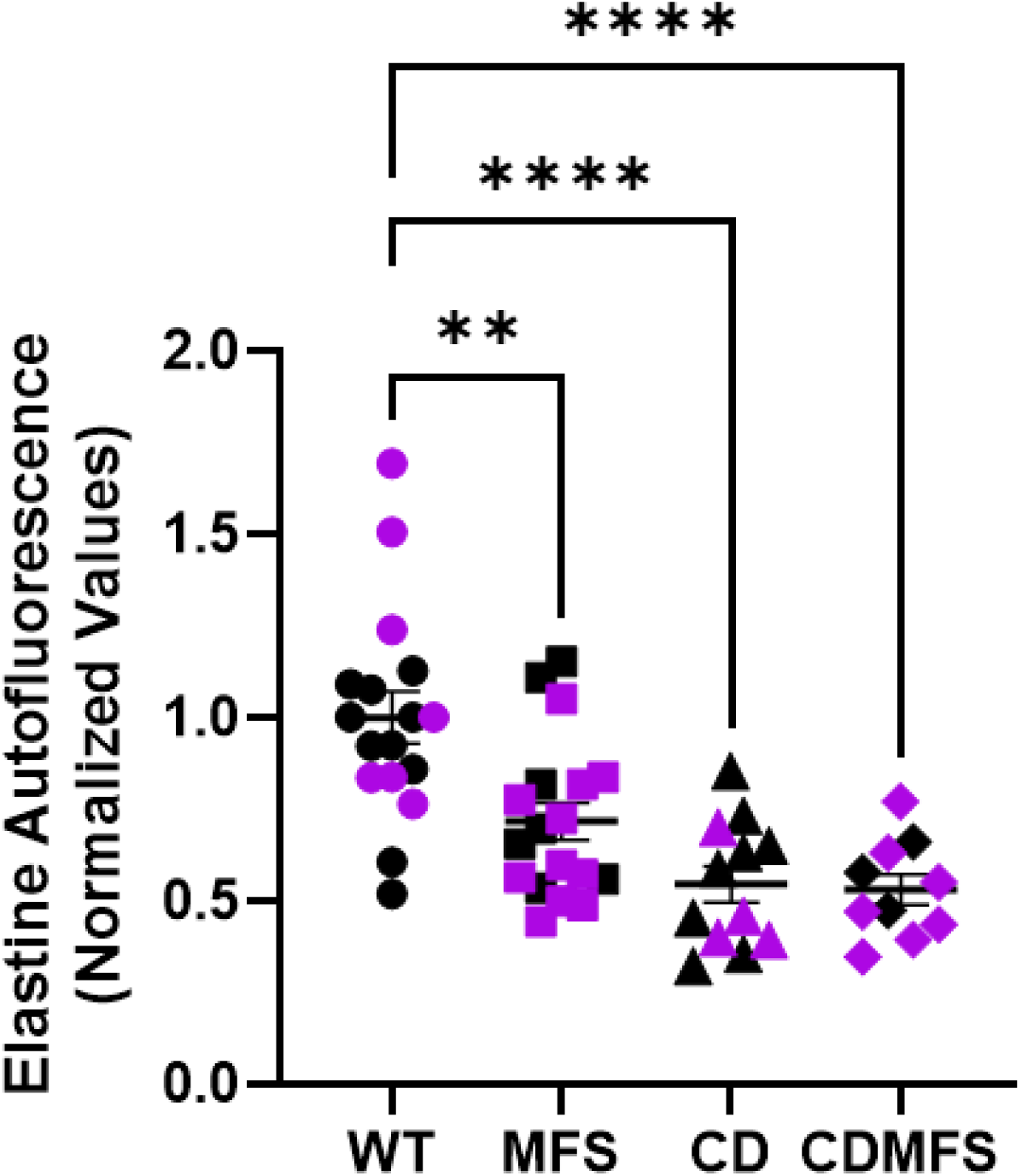
Elastin quantification by autofluorescence. Histogram representing the quantification of elastin autofluorescence in the entire tunica media of the section of the ascending aorta and relativised to the mean of WT littermates. The average per animal was calculated from three different sections. Note that due to ELN haploinsufficiency, CDMFS and CD animals show significantly lower fluorescence emission than WT littermates. The lower fluorescence emission by the media of MFS mice (*p*=0.0012) may be a consequence of the large discontinuities of the elastic laminae. Circles, WT; Squares, MFS; Triangles, CD; Diamonds, CDMFS; Black, males; Purple, females. Data are expressed as the mean ± SEM. Statistical significance was calculated by Dunnet’s comparison test followed by an ordinary one-way ANOVA. \*\**p* < 0.01, \*\*\*\**p* < 0.0001.

**Figure S4.**
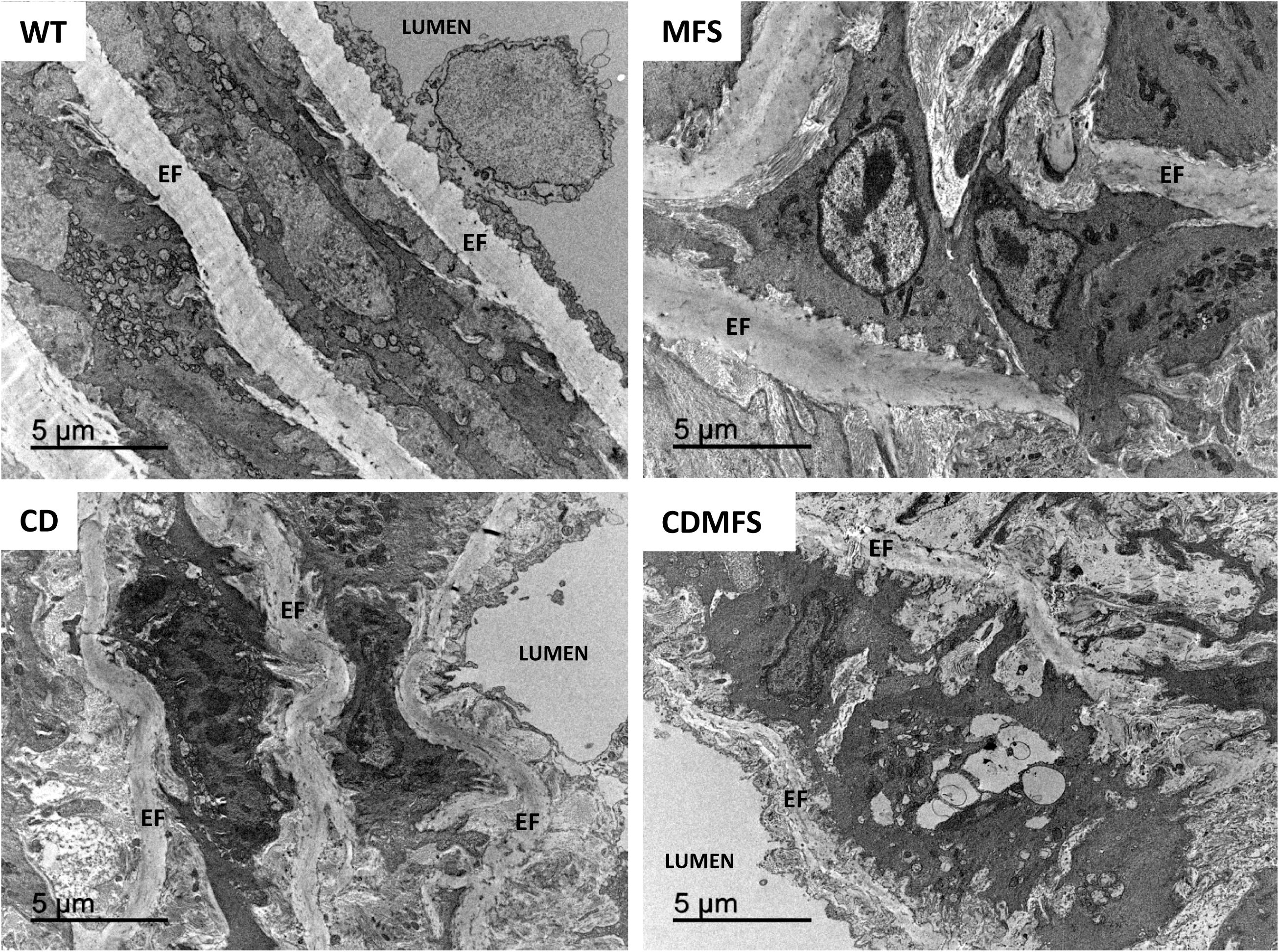
Electron microscopy of the aortic media. Ultrathin sections of WT, MFS, CD and CDMFS aortae display the morphology of aortic media VSMCs and elastic fibres (EL). The MFS image shows a characteristic elastic fibre rupture and disarray of VSMC organisation. CD and CDMFS show thinner elastic fibres as well as altered VSMC morphology. The aorta lumen is indicated. See Results for a more detailed description.

**Figure S5.**
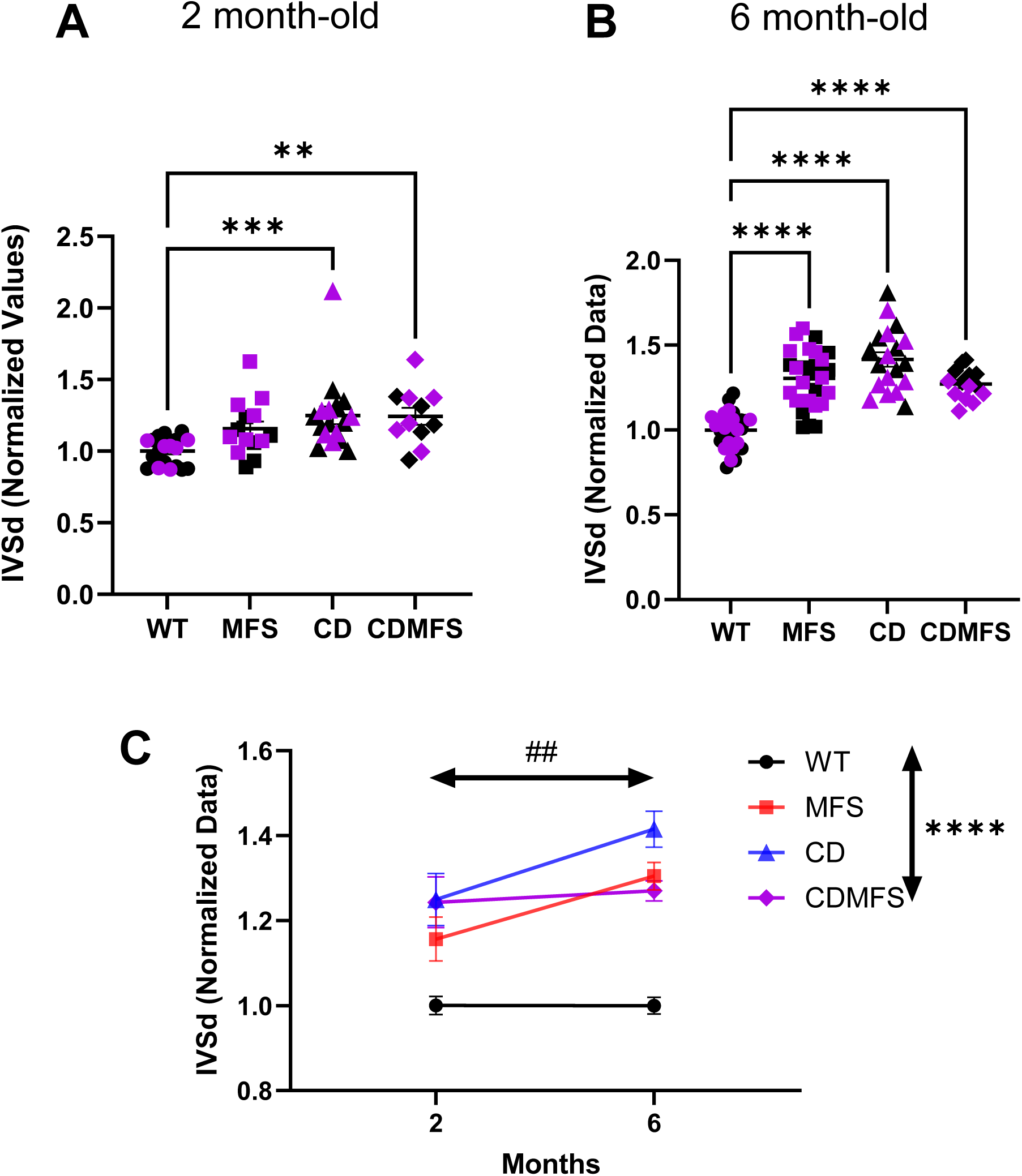
Interventricular septum thickness in diastole (IVSd). Like in CD, the heart of CDMFS mice showed a significant thickening of the IVS both at two (**A**) and six months of age (**B**). At this later age, we observed that MFS animals also presented a significant thickening of this septum (*p*=0.0006), both in males and females. **(C)** Progression of this cardiac parameter over age showing statistical significance between the factors genotype and time. Circles, WT; Squares, MFS; Triangles, CD; Diamonds, CDMFS. Data are expressed as the mean ± SEM. Statistical significance was calculated by Tukey’s comparison test followed by a two-way Mixed effect analysis. *Genotype effect; # time effect. **, ^##^*p* < 0.01; \*\*\**p* < 0.001; \*\*\*\**p* < 0.0001.

**Figure S6.**
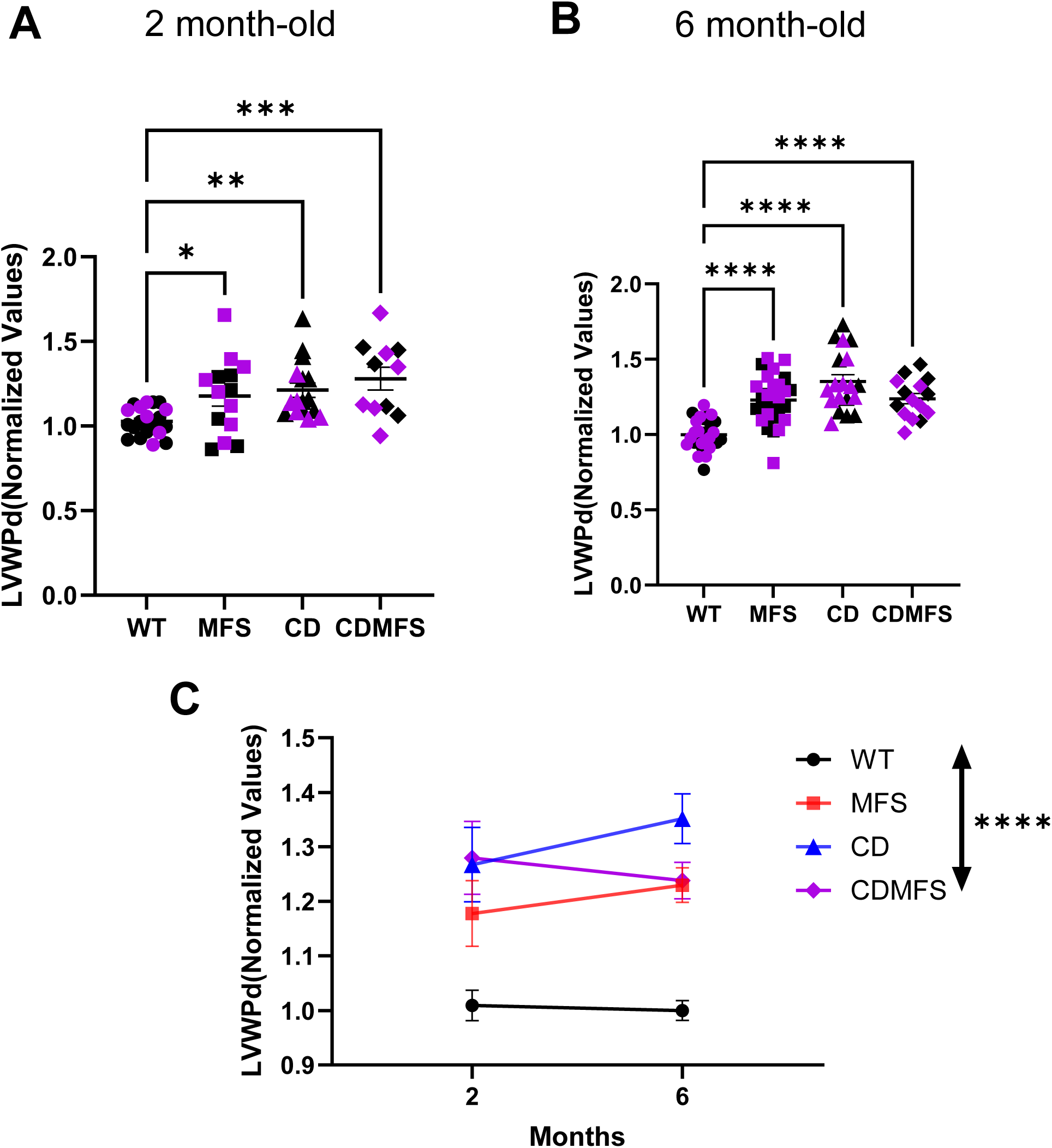
Posterior left ventricular wall thickness in diastole (LVWPd). Unlike the MFS heart, CD and CDMFS mice showed a significant increase in the LVWPd, both at two (**A**) and six months of age (**B**). (**C**) Progression of this cardiac parameter over age showing statistical significance only for the genotype. Circles, WT; Squares, MFS; Triangles, CD; Diamonds, CDMFS. Data are expressed as the mean ± SEM. Statistical significance was calculated by Tukey’s comparison test followed by a two-way Mixed effect analysis. *Genotype effect; \*\**p* < 0.01; \*\*\*\**p* < 0.0001.

**Figure S7.**
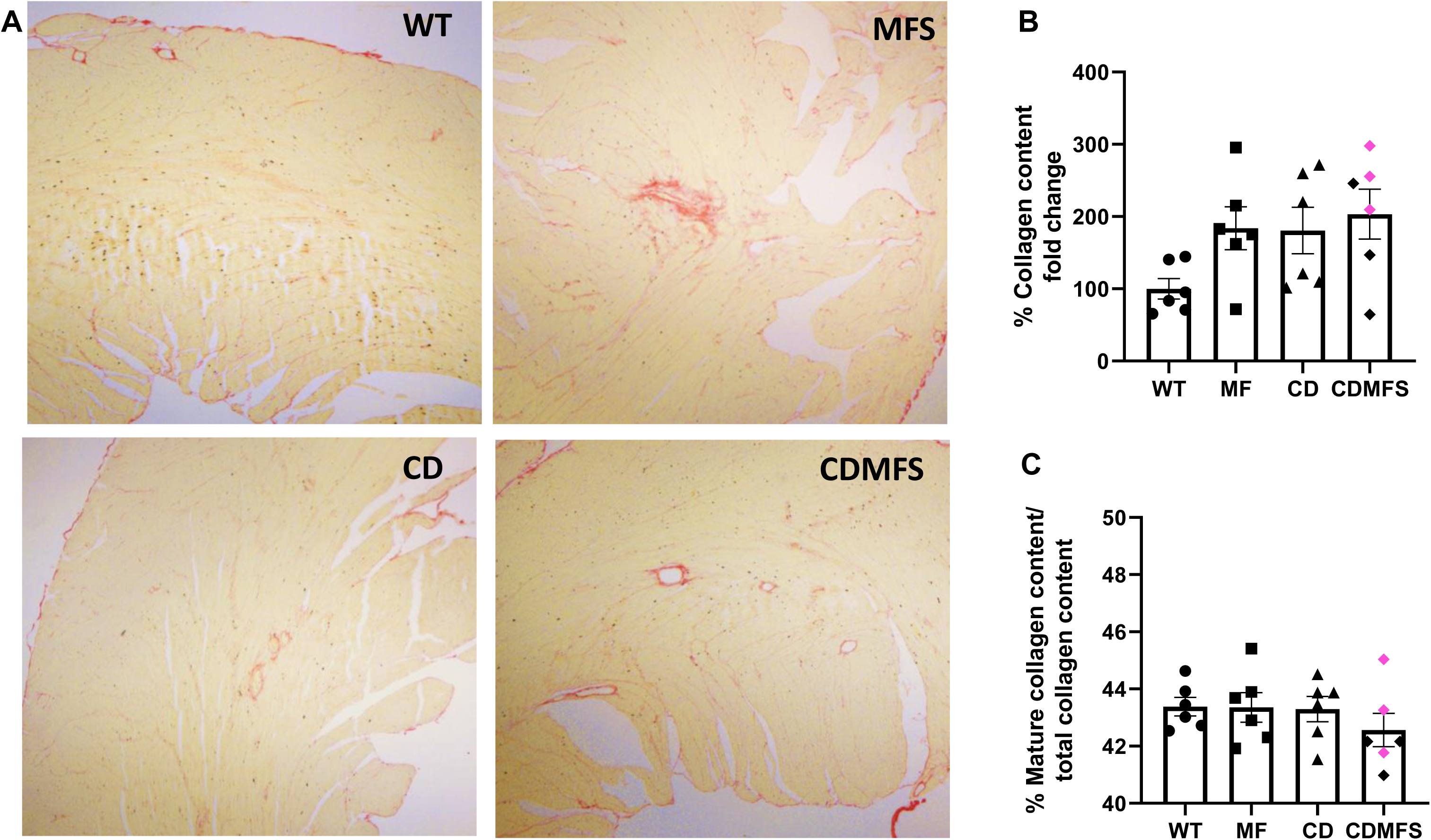
Cardiac fibrosis. Representative histological images of the left ventricle of WT, MFS, CD, and CDMFS hearts stained with Sirius red to reveal collagen fibres **(A)**, and the quantitative analysis of total collagen (**B**; viewed under bright-field microscope) and mature one (**C**; viewed under the polarised light microscope). No significant changes are observed between the genotypes (F_(3,20)=_2.524, *p=*0.0868). Circles, WT; Squares, MFS; Triangles, CD; Diamonds, CDMFS. Data are expressed as the mean ± SEM.

**Figure S8.**
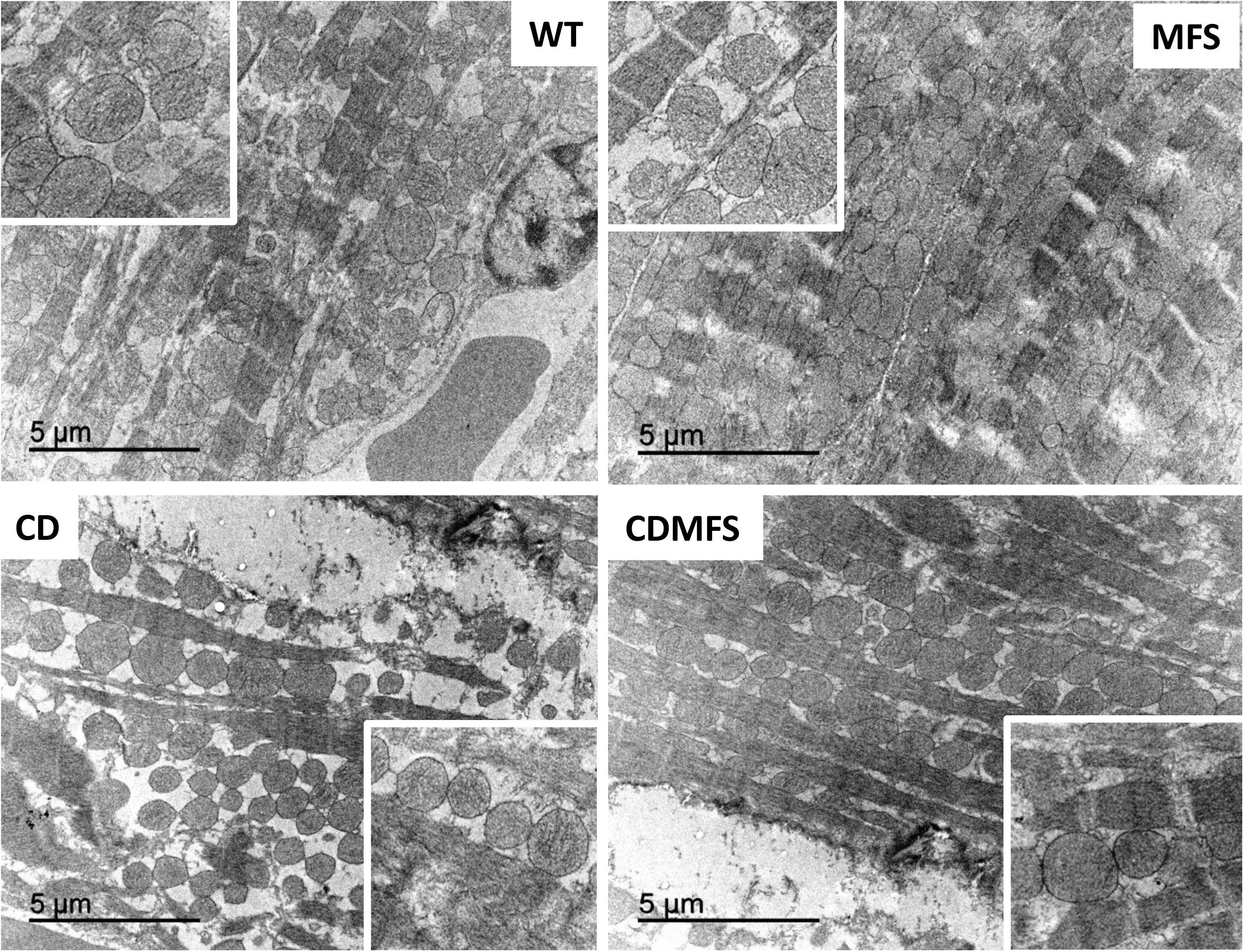
Mitochondrial morphology. Electron microscopy of the cardiac tissue. Ultrathin sections of WT, MFS, CD, and CDMFS heart ventricles showing the morphology and disposition of mitochondria between cardiac fibres. Insets show magnified images of some areas for a better visualisation of mitochondrial morphology.

## SUPPLEMENTAL TABLES

**TABLE S1:** VIABILITY AND BODY WEIGHT.

| Viability |  |  |  |  |  |  |  |  |  |  |  |  |  |  |  |  |
| --- | --- | --- | --- | --- | --- | --- | --- | --- | --- | --- | --- | --- | --- | --- | --- | --- |
| Genotype |  | WT |  |  | MFS |  |  | CD |  |  | CDMFS |  |  |  |  |  |
| Number of pups |  | 41 |  |  | 36 |  |  | 32 |  |  | 21 |  |  |  |  |  |
| Chi-square test |  |  |  |  |  |  |  |  |  |  |  |  |  |  |  |  |
| Chi-square |  |  |  |  | DF |  |  |  |  | P |  |  |  |  |  |  |
| 7.016 |  |  |  |  | 3 |  |  |  |  | 0.0714 |  |  |  |  |  |  |
| Body Weight -Males (g) |  |  |  |  |  |  |  |  |  |  |  |  |  |  |  |  |
| Shapiro-Wilk test | WT-2M | WT-3M | WT-4M | WT-6M | MFS-2M | MFS-3M | MFS-4M | MFS-6M | CD-2M | CD-3M | CD-4M | CD-6M | CDMFS-2M | CDMFS-3M | CDMFS-4M | CDMFS-6M |
| W | 0,9379 | 0,9046 | 0,9212 | 0,9788 | 0,9753 | 0,9579 | 0,8787 | 0,9152 | 0,9721 | 0,9190 | 0,9243 | 0,9436 | 0,8891 | 0,8742 | 0,9083 | 0,9499 |
| P value | 0,3916 | 0,2104 | 0,3285 | 0,9606 | 0,9148 | 0,7455 | 0,1000 | 0,2159 | 0,9074 | 0,3836 | 0,4295 | 0,5935 | 0,3789 | 0,2837 | 0,4572 | 0,7398 |
| ONE- WAY ANOVA-2M |  |  |  |  |  |  |  |  |  |  |  |  |  |  |  |  |
| ANOVA table |  |  |  |  | F (DFn, DFd) |  |  |  |  | P value |  |  |  |  |  |  |
| Genotype |  |  |  |  | F (3, 41) = 16,16 |  |  |  |  | P<0,0001 |  |  |  |  |  |  |
| Dunnett's multiple comparisons test |  |  |  |  |  |  |  |  | Adjusted P Value |  |  |  |  |  |  |  |
| WT vs. MFS |  |  |  |  |  |  |  |  | 0,8306 |  |  |  |  |  |  |  |
| WT vs. CD |  |  |  |  |  |  |  |  | 0,0001 |  |  |  |  |  |  |  |
| WT vs. CDMFS |  |  |  |  |  |  |  |  | 0,0002 |  |  |  |  |  |  |  |
| ONE- WAY ANOVA-6M |  |  |  |  |  |  |  |  |  |  |  |  |  |  |  |  |
| ANOVA table |  |  |  |  | F (DFn, DFd) |  |  |  |  | P value |  |  |  |  |  |  |
| Genotype |  |  |  |  | F (3, 40) = 11,39 |  |  |  |  | P<0,0001 |  |  |  |  |  |  |
| Dunnett's multiple comparisons test |  |  |  |  |  |  |  |  | Adjusted P Value |  |  |  |  |  |  |  |
| WT vs. MFS |  |  |  |  |  |  |  |  | 0,9997 |  |  |  |  |  |  |  |
| WT vs. CD |  |  |  |  |  |  |  |  | 0,0001 |  |  |  |  |  |  |  |
| WT vs. CDMFS |  |  |  |  |  |  |  |  | 0,0026 |  |  |  |  |  |  |  |
| PROGRESSION |  |  |  |  |  |  |  |  |  |  |  |  |  |  |  |  |
| Mixed-effects model (REML) |  |  |  |  | F (DFn, DFd) |  |  |  |  | P value |  |  |  |  |  |  |
| Time |  |  |  |  | F (1,847, 62,17) = 106,4 |  |  |  |  | <0,0001 |  |  |  |  |  |  |
| GENOTYPE |  |  |  |  | F (3, 44) = 53,28 |  |  |  |  | <0,0001 |  |  |  |  |  |  |
| Time x GENOTYPE |  |  |  |  | F (9, 101) = 0,8470 |  |  |  |  | 0,5750 |  |  |  |  |  |  |

| Body Weight -Females |  |  |  |  |  |  |  |  |  |  |  |  |  |  |  |  |
| --- | --- | --- | --- | --- | --- | --- | --- | --- | --- | --- | --- | --- | --- | --- | --- | --- |
| Shapiro-Wilk test | WT-2M | WT-3M | WT-4M | WT-6M | MFS-2M | MFS-3M | MFS-4M | MFS-6M | CD-2M | CD-3M | CD-4M | CD-6M | CDMFS-2M | CDMFS-3M | CDMFS-4M | CDMFS-6M |
| W | 0,9661 | 0,9066 | 0,9077 | 0,9671 | 0,9074 | 0,9387 | 0,9402 | 0,9617 | 0,9370 | 0,8769 | 0,9614 | 0,9227 | 0,9863 | 0,9250 | 0,9182 | 0,9416 |
| P value | 0,8213 | 0,2224 | 0,0783 | 0,7898 | 0,2636 | 0,6271 | 0,6135 | 0,8084 | 0,5512 | 0,0800 | 0,7891 | 0,4154 | 0,9843 | 0,5089 | 0,4557 | 0,5984 |
| ONE- WAY ANOVA-2M |  |  |  |  |  |  |  |  |  |  |  |  |  |  |  |  |
| ANOVA table |  |  |  | F (DFn, DFd) |  |  |  |  |  |  |  | P value |  |  |  |  |
| Genotype |  |  |  | F (3, 36) = 12,51 |  |  |  |  |  |  |  | P<0,0001 |  |  |  |  |
| Dunnett's multiple comparisons test |  |  |  |  |  |  |  | Adjusted P Value |  |  |  |  |  |  |  |  |
| WT vs. MFS |  |  |  |  |  |  |  | 0,4724 |  |  |  |  |  |  |  |  |
| WT vs. CD |  |  |  |  |  |  |  | 0,0022 |  |  |  |  |  |  |  |  |
| WT vs. CDMFS |  |  |  |  |  |  |  | 0,0008 |  |  |  |  |  |  |  |  |
| ONE- WAY ANOVA-6M |  |  |  |  |  |  |  |  |  |  |  |  |  |  |  |  |
| ANOVA table |  |  |  | F (DFn, DFd) |  |  |  |  |  |  |  | P value |  |  |  |  |
| Genotype |  |  |  | F (3, 42) = 8,883 |  |  |  |  |  |  |  | P=0,0001 |  |  |  |  |
| Dunnett's multiple comparisons test |  |  |  |  |  |  |  | Adjusted P Value |  |  |  |  |  |  |  |  |
| WT vs. MFS |  |  |  |  |  |  |  | 0,3771 |  |  |  |  |  |  |  |  |
| WT vs. CD |  |  |  |  |  |  |  | <0,0001 |  |  |  |  |  |  |  |  |
| WT vs. CDMFS |  |  |  |  |  |  |  | 0,0044 |  |  |  |  |  |  |  |  |
| PROGRESSION |  |  |  |  |  |  |  |  |  |  |  |  |  |  |  |  |
| Mixed-effects model (REML) |  |  |  | F (DFn, DFd) |  |  |  |  |  |  |  | P value |  |  |  |  |
| Time |  |  |  | F (2,142, 74,27) = 98,19 |  |  |  |  |  |  |  | <0,0001 |  |  |  |  |
| GENOTYPE |  |  |  | F (3, 47) = 24,67 |  |  |  |  |  |  |  | <0,0001 |  |  |  |  |
| Time x GENOTYPE |  |  |  | F (9, 104) = 1,094 |  |  |  |  |  |  |  | 0,3741 |  |  |  |  |

**TABLE S2:**
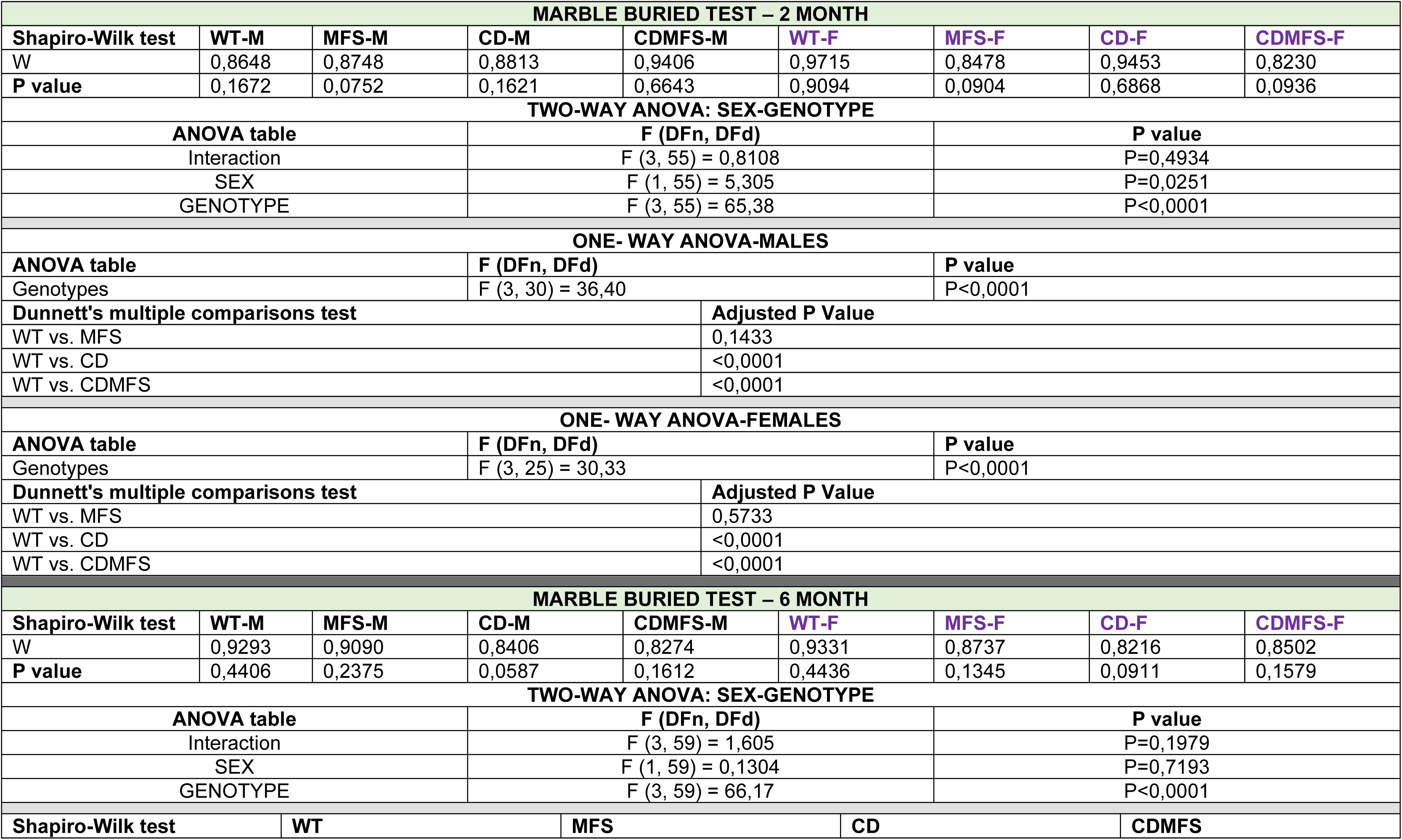
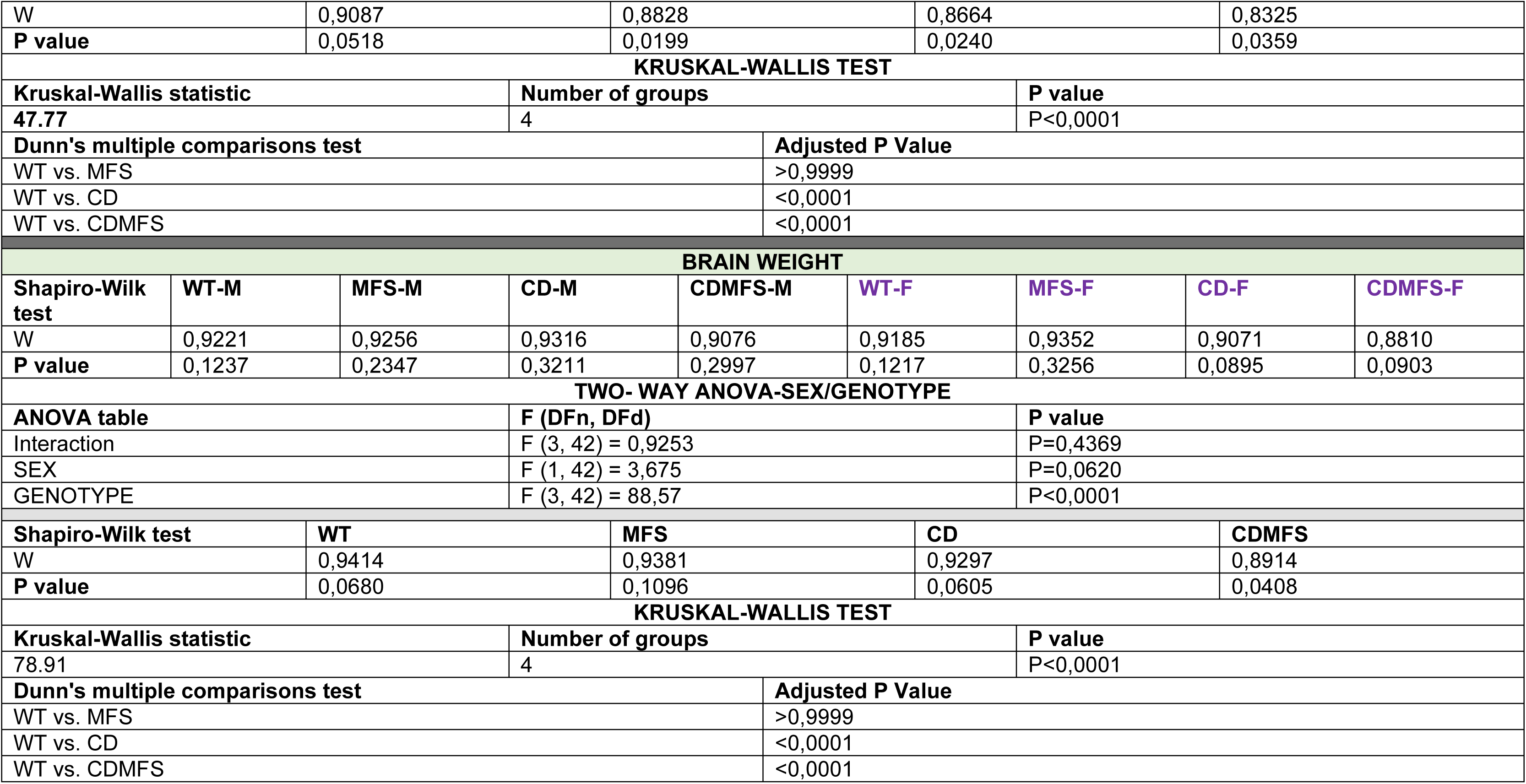
BEHAVIOR-ASSOCIATED ALTERATIONS AND BRAIN WEIGHT.

**TABLE S3:**
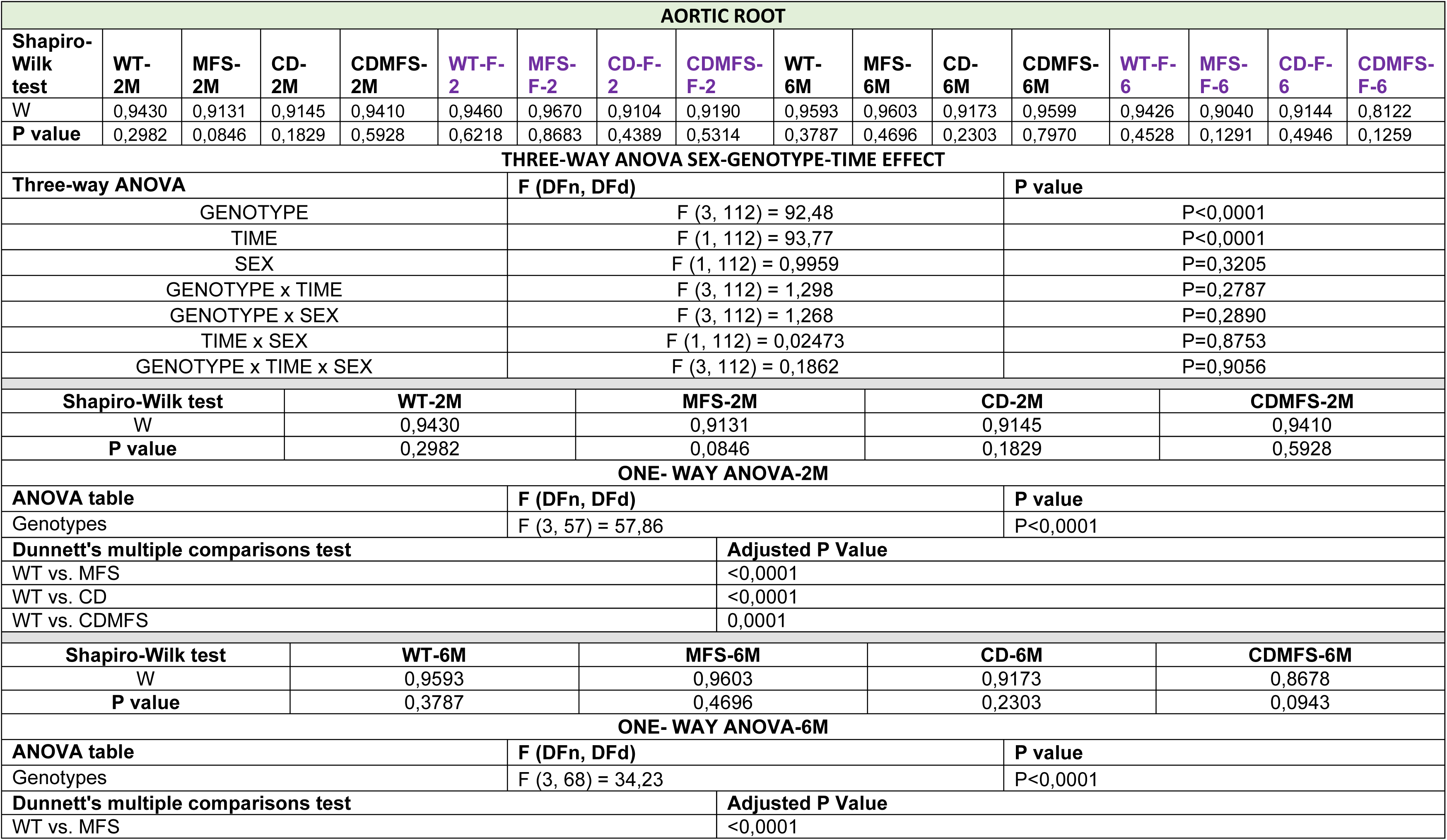
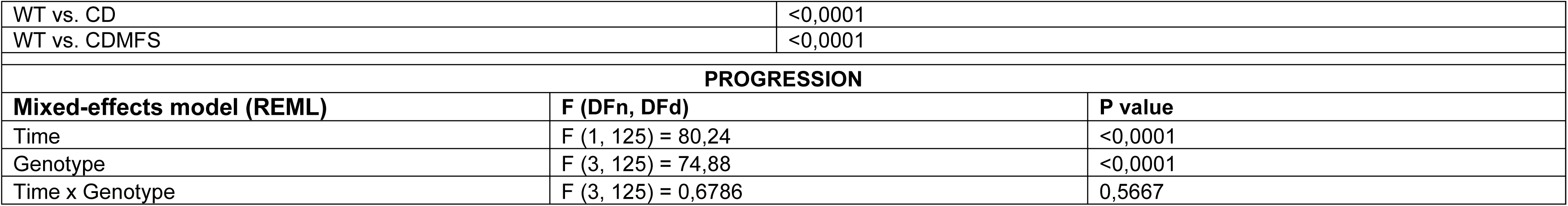
AORTIC ROOT.

| AORTIC ROOT |  |  |  |  |  |  |  |  |  |  |  |  |  |  |  |  |
| --- | --- | --- | --- | --- | --- | --- | --- | --- | --- | --- | --- | --- | --- | --- | --- | --- |
| Shapiro-Wilk test | WT-2M | MFS-2M | CD-2M | CDMFS-2M | WT-F-2 | MFS-F-2 | CD-F-2 | CDMFS-F-2 | WT-6M | MFS-6M | CD-6M | CDMFS-6M | WT-F-6 | MFS-F-6 | CD-F-6 | CDMFS-F-6 |
| W | 0,9430 | 0,9131 | 0,9145 | 0,9410 | 0,9460 | 0,9670 | 0,9104 | 0,9190 | 0,9593 | 0,9603 | 0,9173 | 0,9599 | 0,9426 | 0,9040 | 0,9144 | 0,8122 |
| P value | 0,2982 | 0,0846 | 0,1829 | 0,5928 | 0,6218 | 0,8683 | 0,4389 | 0,5314 | 0,3787 | 0,4696 | 0,2303 | 0,7970 | 0,4528 | 0,1291 | 0,4946 | 0,1259 |
| THREE-WAY ANOVA SEX-GENOTYPE-TIME EFFECT |  |  |  |  |  |  |  |  |  |  |  |  |  |  |  |  |
| Three-way ANOVA |  |  |  |  | F (DFn, DFd) |  |  |  |  | P value |  |  |  |  |  |  |
| GENOTYPE |  |  |  |  | F (3, 112) = 92,48 |  |  |  |  | P<0,0001 |  |  |  |  |  |  |
| TIME |  |  |  |  | F (1, 112) = 93,77 |  |  |  |  | P<0,0001 |  |  |  |  |  |  |
| SEX |  |  |  |  | F (1, 112) = 0,9959 |  |  |  |  | P=0,3205 |  |  |  |  |  |  |
| GENOTYPE x TIME |  |  |  |  | F (3, 112) = 1,298 |  |  |  |  | P=0,2787 |  |  |  |  |  |  |
| GENOTYPE x SEX |  |  |  |  | F (3, 112) = 1,268 |  |  |  |  | P=0,2890 |  |  |  |  |  |  |
| TIME x SEX |  |  |  |  | F (1, 112) = 0,02473 |  |  |  |  | P=0,8753 |  |  |  |  |  |  |
| GENOTYPE x TIME x SEX |  |  |  |  | F (3, 112) = 0,1862 |  |  |  |  | P=0,9056 |  |  |  |  |  |  |
| Shapiro-Wilk test |  |  | WT-2M |  |  | MFS-2M |  |  | CD-2M |  |  | CDMFS-2M |  |  |  |  |
| W |  |  | 0,9430 |  |  | 0,9131 |  |  | 0,9145 |  |  | 0,9410 |  |  |  |  |
| P value |  |  | 0,2982 |  |  | 0,0846 |  |  | 0,1829 |  |  | 0,5928 |  |  |  |  |
| ONE- WAY ANOVA-2M |  |  |  |  |  |  |  |  |  |  |  |  |  |  |  |  |
| ANOVA table |  |  |  |  | F (DFn, DFd) |  |  |  |  | P value |  |  |  |  |  |  |
| Genotypes |  |  |  |  | F (3, 57) = 57,86 |  |  |  |  | P<0,0001 |  |  |  |  |  |  |
| Dunnett's multiple comparisons test |  |  |  |  |  |  |  |  | Adjusted P Value |  |  |  |  |  |  |  |
| WT vs. MFS |  |  |  |  |  |  |  |  | <0,0001 |  |  |  |  |  |  |  |
| WT vs. CD |  |  |  |  |  |  |  |  | <0,0001 |  |  |  |  |  |  |  |
| WT vs. CDMFS |  |  |  |  |  |  |  |  | 0,0001 |  |  |  |  |  |  |  |
| Shapiro-Wilk test |  |  | WT-6M |  |  | MFS-6M |  |  | CD-6M |  |  | CDMFS-6M |  |  |  |  |
| W |  |  | 0,9593 |  |  | 0,9603 |  |  | 0,9173 |  |  | 0,8678 |  |  |  |  |
| P value |  |  | 0,3787 |  |  | 0,4696 |  |  | 0,2303 |  |  | 0,0943 |  |  |  |  |
| ONE- WAY ANOVA-6M |  |  |  |  |  |  |  |  |  |  |  |  |  |  |  |  |
| ANOVA table |  |  |  |  | F (DFn, DFd) |  |  |  |  | P value |  |  |  |  |  |  |
| Genotypes |  |  |  |  | F (3, 68) = 34,23 |  |  |  |  | P<0,0001 |  |  |  |  |  |  |
| Dunnett's multiple comparisons test |  |  |  |  |  |  |  |  | Adjusted P Value |  |  |  |  |  |  |  |
| WT vs. MFS |  |  |  |  |  |  |  |  | <0,0001 |  |  |  |  |  |  |  |

|  |  |  |
| --- | --- | --- |
| WT vs. CD | <0,0001 |  |
| WT vs. CDMFS | <0,0001 |  |
| PROGRESSION |  |  |
| Mixed-effects model (REML) | F (DFn, DFd) | P value |
| Time | F (1, 125) = 80,24 | <0,0001 |
| Genotype | F (3, 125) = 74,88 | <0,0001 |
| Time x Genotype | F (3, 125) = 0,6786 | 0,5667 |

**TABLE S4:**
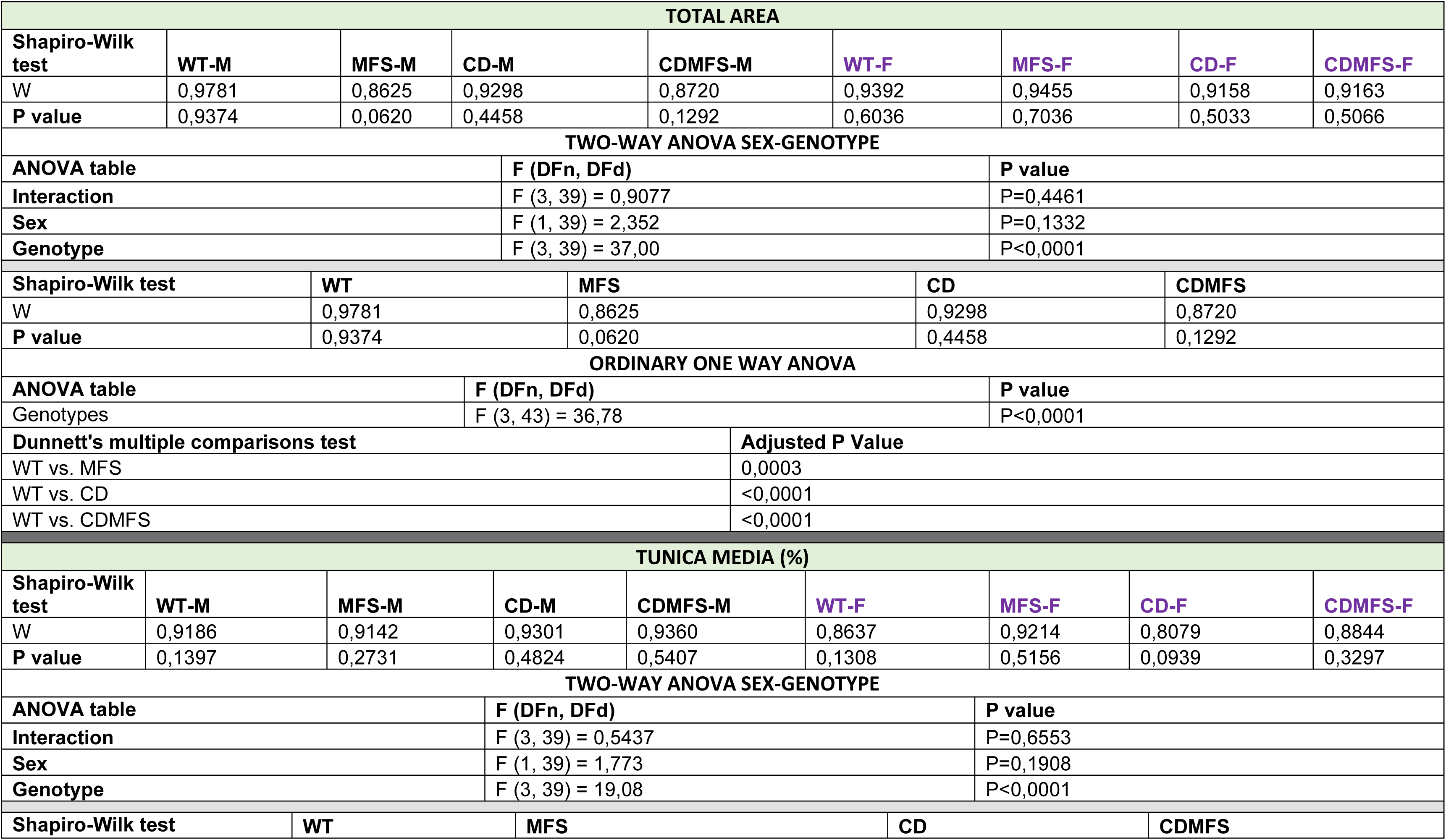
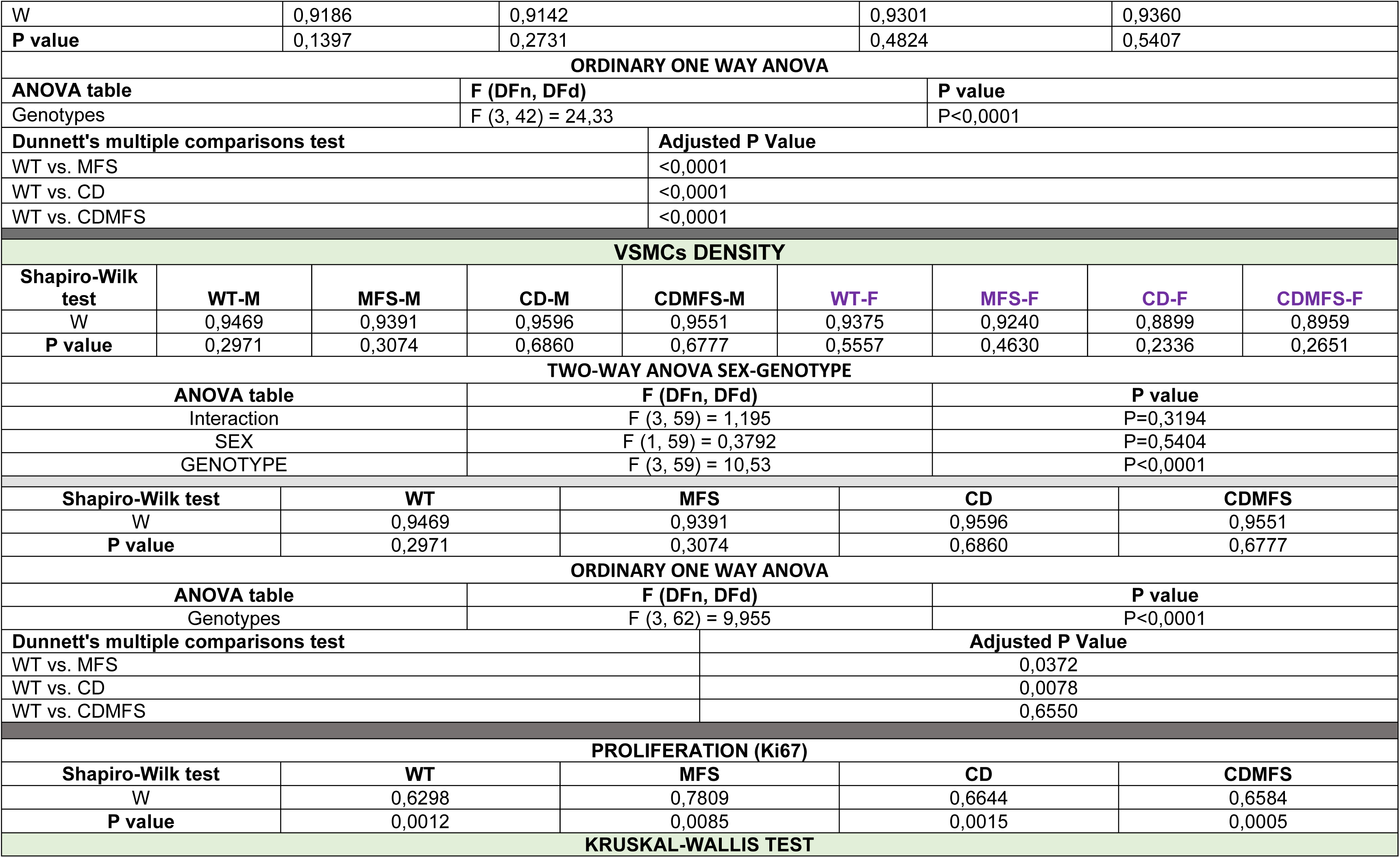
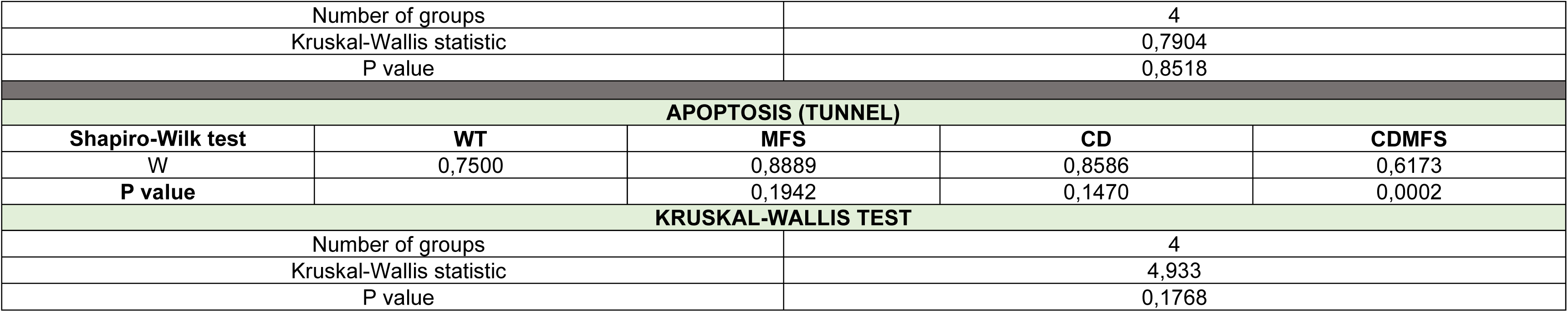
HISTOLOGICAL OUTCOMES.

**TABLE S5:** ELASTIC LAMINAE INTEGRITY.

| LARGE BREAKS |  |  |  |  |  |  |  |  |
| --- | --- | --- | --- | --- | --- | --- | --- | --- |
| Shapiro-Wilk test | WT-M | MFS-M | CD-M | CDMFS-M | WT-F | MFS-F | CD-F | CDMFS-F |
| W | 0,7875 | 0,9132 | 0,8944 | 0,8527 | 0,8455 | 0,9112 | 0,9714 | 0,8950 |
| P value | 0,0006 | 0,2026 | 0,1577 | 0,0309 | 0,0513 | 0,4747 | 0,8500 | 0,2601 |
| Kruskal-Wallis test |  |  |  |  |  |  |  |  |
| Kruskal-Wallis statistic |  |  |  | 46.11 |  |  |  |  |
| Number of groups |  |  |  | 8 |  |  |  |  |
| P value |  |  |  | P<0,0001 |  |  |  |  |
| Dunn's multiple comparisons test |  |  |  | Adjusted P Value |  |  |  |  |
| WT-Male vs. WT-Female |  |  |  | >0,9999 |  |  |  |  |
| MFS-Male vs. MFS-Female |  |  |  | >0,9999 |  |  |  |  |
| CD-Male vs. CD-Female |  |  |  | >0,9999 |  |  |  |  |
| CDMFS-Male vs. CDMFS-Female |  |  |  | >0,9999 |  |  |  |  |
| Shapiro-Wilk test |  | WT |  | MFS |  | CD |  | CDMFS |
| W |  | 0,7875 |  | 0,9132 |  | 0,8944 |  | 0,8527 |
| P value |  | 0,0006 |  | 0,2026 |  | 0,1577 |  | 0,0309 |
| Kruskal-Wallis test |  |  |  |  |  |  |  |  |
| Kruskal-Wallis statistic |  |  |  | 34.09 |  |  |  |  |
| Number of groups |  |  |  | 4 |  |  |  |  |
| P value |  |  |  | P<0,0001 |  |  |  |  |
| Dunn's multiple comparisons test |  |  |  | Adjusted P Value |  |  |  |  |
| WT vs. MFS |  |  |  | <0,0001 |  |  |  |  |
| WT vs. CD |  |  |  | 0,3907 |  |  |  |  |
| WT vs. CDMFS |  |  |  | 0,0014 |  |  |  |  |
| UNDULATION FACTOR |  |  |  |  |  |  |  |  |
| Shapiro-Wilk test | WT-M | MFS-M | CD-M | CDMFS-M | WT-F | MFS-F | CD-F | CDMFS-F |
| W | 0,8947 | 0,7861 | 0,9044 | 0,9537 | 0,8888 | 0,9293 | 0,8528 | 0,9037 |
| P value | 0,1355 | 0,0203 | 0,3583 | 0,7638 | 0,1939 | 0,5451 | 0,2481 | 0,3541 |
| Kruskal-Wallis test |  |  |  |  |  |  |  |  |
| Kruskal-Wallis statistic |  |  |  | 35.02 |  |  |  |  |
| Number of groups |  |  |  | 8 |  |  |  |  |
| P value |  |  |  | P<0,0001 |  |  |  |  |
| Dunn's multiple comparisons test |  |  |  | Adjusted P Value |  |  |  |  |
| WT-Male vs. WT-Female |  |  |  | >0,9999 |  |  |  |  |
| MFS-Male vs. MFS-Female |  |  |  | >0,9999 |  |  |  |  |
| CD-Male vs. CD-Female |  |  |  | 0,5606 |  |  |  |  |
| CDMFS-Male vs. CDMFS-Female |  |  |  | >0,9999 |  |  |  |  |
| Shapiro-Wilk test |  | WT |  | MFS |  | CD |  | CDMFS |
| W |  | 0,9779 |  | 0,8913 |  | 0,9287 |  | 0,9578 |
| P value |  | 0,8925 |  | 0,0702 |  | 0,4355 |  | 0,7523 |
| ORDINARY ONE WAY ANOVA |  |  |  |  |  |  |  |  |
| ANOVA table |  |  | F (DFn, DFd) |  |  | P value |  |  |
| Genotypes |  |  | F (3, 54) = 23,67 |  |  | P<0,0001 |  |  |
| Dunnnett's multiple comparisons test |  |  |  |  | Adjusted P Value |  |  |  |
| WT vs. MFS |  |  |  |  | <0,0001 |  |  |  |
| WT vs. CD |  |  |  |  | 0,9966 |  |  |  |
| WT vs. CDMFS |  |  |  |  | <0,0001 |  |  |  |
| ELN AUTOFLUORESCENCE |  |  |  |  |  |  |  |  |
| Shapiro-Wilk test | WT-M | MFS-M | CD-M | CDMFS-M | WT-F | MFS-F | CD-F | CDMFS-F |
| W | 0,9409 | 0,9074 | 0,9387 | 0,9728 | 0,8873 | 0,9279 | 0,7775 | 0,9449 |
| P value | 0,3288 | 0,0776 | 0,4809 | 0,9153 | 0,2609 | 0,3901 | 0,0676 | 0,6836 |
| TWO-WAY ANOVA SEX-GENOTYPE |  |  |  |  |  |  |  |  |
| ANOVA table |  |  | F (DFn, DFd) |  |  | P value |  |  |
| Interaction |  |  | F (3, 49) = 1,904 |  |  | P=0,1412 |  |  |
| SEX |  |  | F (1, 49) = 0,03813 |  |  | P=0,8460 |  |  |
| GENOTYPE |  |  | F (3, 49) = 14,43 |  |  | P<0,0001 |  |  |
| Shapiro-Wilk test |  | WT |  | MFS |  | CD |  | CDMFS |
| W |  | 0,9409 |  | 0,9074 |  | 0,9387 |  | 0,9728 |
| P value |  | 0,3288 |  | 0,0776 |  | 0,4809 |  | 0,9153 |
| ORDINARY ONE WAY ANOVA |  |  |  |  |  |  |  |  |
| ANOVA table |  |  | F (DFn, DFd) |  |  | P value |  |  |
| Genotypes |  |  | F (3, 53) = 13,83 |  |  | P<0,0001 |  |  |
| Dunnnett's multiple comparisons test |  |  |  |  | Adjusted P Value |  |  |  |
| WT vs. MFS |  |  |  |  | 0,0012 |  |  |  |
| WT vs. CD |  |  |  |  | <0,0001 |  |  |  |
| WT vs. CDMFS |  |  |  |  | <0,0001 |  |  |  |

**TABLE S6:**
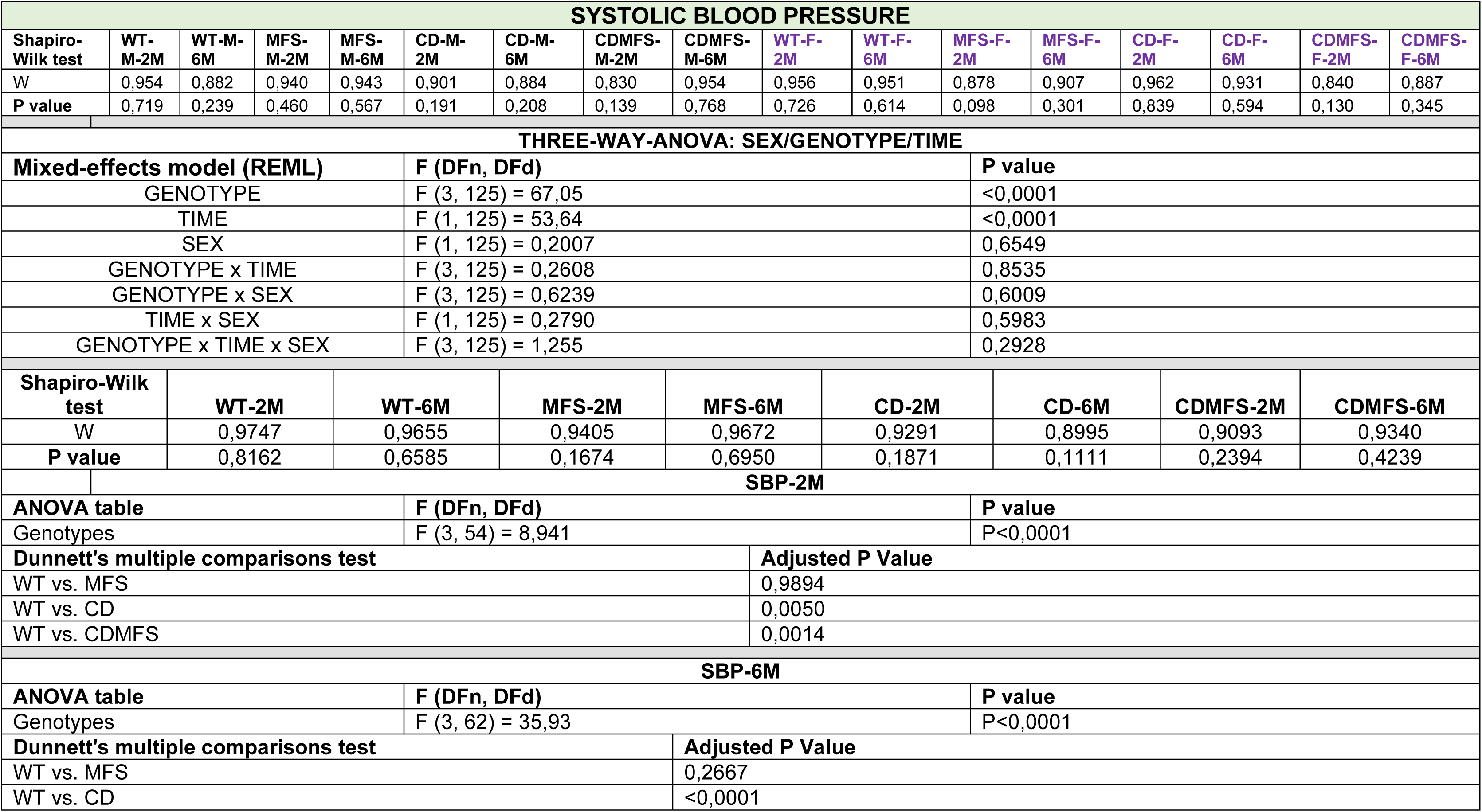
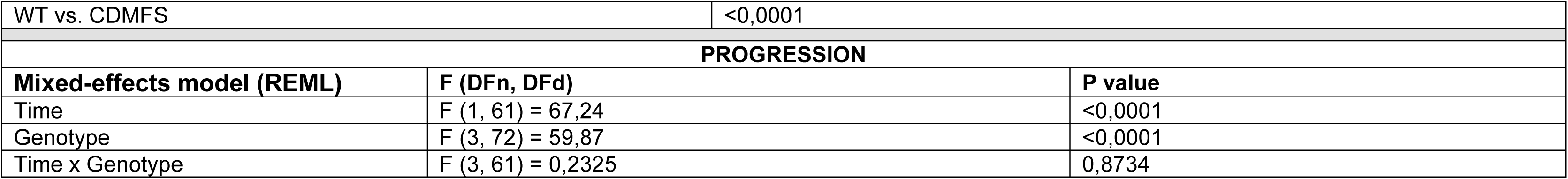
SYSTOLIC BLOOD PRESSURE.

| SYSTOLIC BLOOD PRESSURE |  |  |  |  |  |  |  |  |  |  |  |  |  |  |  |  |
| --- | --- | --- | --- | --- | --- | --- | --- | --- | --- | --- | --- | --- | --- | --- | --- | --- |
| Shapiro-Wilk test | WT-M-2M | WT-M-6M | MFS-M-2M | MFS-M-6M | CD-M-2M | CD-M-6M | CDMFS-M-2M | CDMFS-M-6M | WT-F-2M | WT-F-6M | MFS-F-2M | MFS-F-6M | CD-F-2M | CD-F-6M | CDMFS-F-2M | CDMFS-F-6M |
| W | 0,954 | 0,882 | 0,940 | 0,943 | 0,901 | 0,884 | 0,830 | 0,954 | 0,956 | 0,951 | 0,878 | 0,907 | 0,962 | 0,931 | 0,840 | 0,887 |
| P value | 0,719 | 0,239 | 0,460 | 0,567 | 0,191 | 0,208 | 0,139 | 0,768 | 0,726 | 0,614 | 0,098 | 0,301 | 0,839 | 0,594 | 0,130 | 0,345 |
| THREE-WAY-ANOVA: SEX/GENOTYPE/TIME |  |  |  |  |  |  |  |  |  |  |  |  |  |  |  |  |
| Mixed-effects model (REML) |  |  |  |  | F (DFn, DFd) |  |  |  |  | P value |  |  |  |  |  |  |
| GENOTYPE |  |  |  |  | F (3, 125) = 67,05 |  |  |  |  | <0,0001 |  |  |  |  |  |  |
| TIME |  |  |  |  | F (1, 125) = 53,64 |  |  |  |  | <0,0001 |  |  |  |  |  |  |
| SEX |  |  |  |  | F (1, 125) = 0,2007 |  |  |  |  | 0,6549 |  |  |  |  |  |  |
| GENOTYPE x TIME |  |  |  |  | F (3, 125) = 0,2608 |  |  |  |  | 0,8535 |  |  |  |  |  |  |
| GENOTYPE x SEX |  |  |  |  | F (3, 125) = 0,6239 |  |  |  |  | 0,6009 |  |  |  |  |  |  |
| TIME x SEX |  |  |  |  | F (1, 125) = 0,2790 |  |  |  |  | 0,5983 |  |  |  |  |  |  |
| GENOTYPE x TIME x SEX |  |  |  |  | F (3, 125) = 1,255 |  |  |  |  | 0,2928 |  |  |  |  |  |  |
| Shapiro-Wilk test |  | WT-2M |  | WT-6M |  | MFS-2M |  | MFS-6M |  | CD-2M |  | CD-6M |  | CDMFS-2M |  | CDMFS-6M |
| W |  | 0,9747 |  | 0,9655 |  | 0,9405 |  | 0,9672 |  | 0,9291 |  | 0,8995 |  | 0,9093 |  | 0,9340 |
| P value |  | 0,8162 |  | 0,6585 |  | 0,1674 |  | 0,6950 |  | 0,1871 |  | 0,1111 |  | 0,2394 |  | 0,4239 |
| SBP-2M |  |  |  |  |  |  |  |  |  |  |  |  |  |  |  |  |
| ANOVA table |  |  |  |  | F (DFn, DFd) |  |  |  |  | P value |  |  |  |  |  |  |
| Genotypes |  |  |  |  | F (3, 54) = 8,941 |  |  |  |  | P<0,0001 |  |  |  |  |  |  |
| Dunnett's multiple comparisons test |  |  |  |  |  |  |  |  | Adjusted P Value |  |  |  |  |  |  |  |
| WT vs. MFS |  |  |  |  |  |  |  |  | 0,9894 |  |  |  |  |  |  |  |
| WT vs. CD |  |  |  |  |  |  |  |  | 0,0050 |  |  |  |  |  |  |  |
| WT vs. CDMFS |  |  |  |  |  |  |  |  | 0,0014 |  |  |  |  |  |  |  |
| SBP-6M |  |  |  |  |  |  |  |  |  |  |  |  |  |  |  |  |
| ANOVA table |  |  |  |  | F (DFn, DFd) |  |  |  |  | P value |  |  |  |  |  |  |
| Genotypes |  |  |  |  | F (3, 62) = 35,93 |  |  |  |  | P<0,0001 |  |  |  |  |  |  |
| Dunnett's multiple comparisons test |  |  |  |  |  |  |  |  | Adjusted P Value |  |  |  |  |  |  |  |
| WT vs. MFS |  |  |  |  |  |  |  |  | 0,2667 |  |  |  |  |  |  |  |
| WT vs. CD |  |  |  |  |  |  |  |  | <0,0001 |  |  |  |  |  |  |  |

|  |  |  |
| --- | --- | --- |
| WT vs. CDMFS |  | <0,0001 |
| **PROGRESSION** |  |  |
| **Mixed-effects model (REML)** | **F (DFn, DFd)** | **P value** |
| Time | F (1, 61) = 67,24 | <0,0001 |
| Genotype | F (3, 72) = 59,87 | <0,0001 |
| Time x Genotype | F (3, 61) = 0,2325 | 0,8734 |

**TABLE S7:**
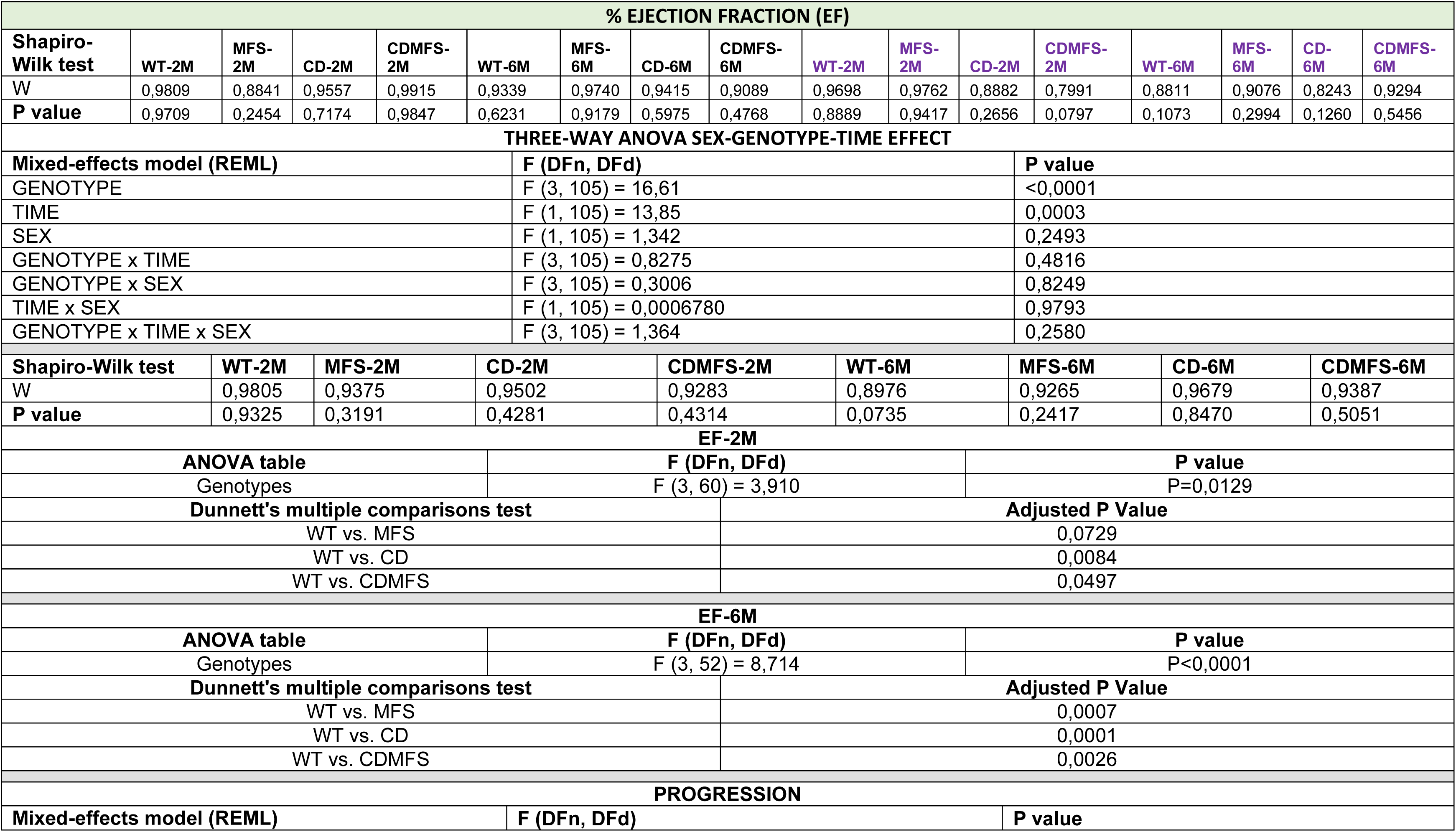
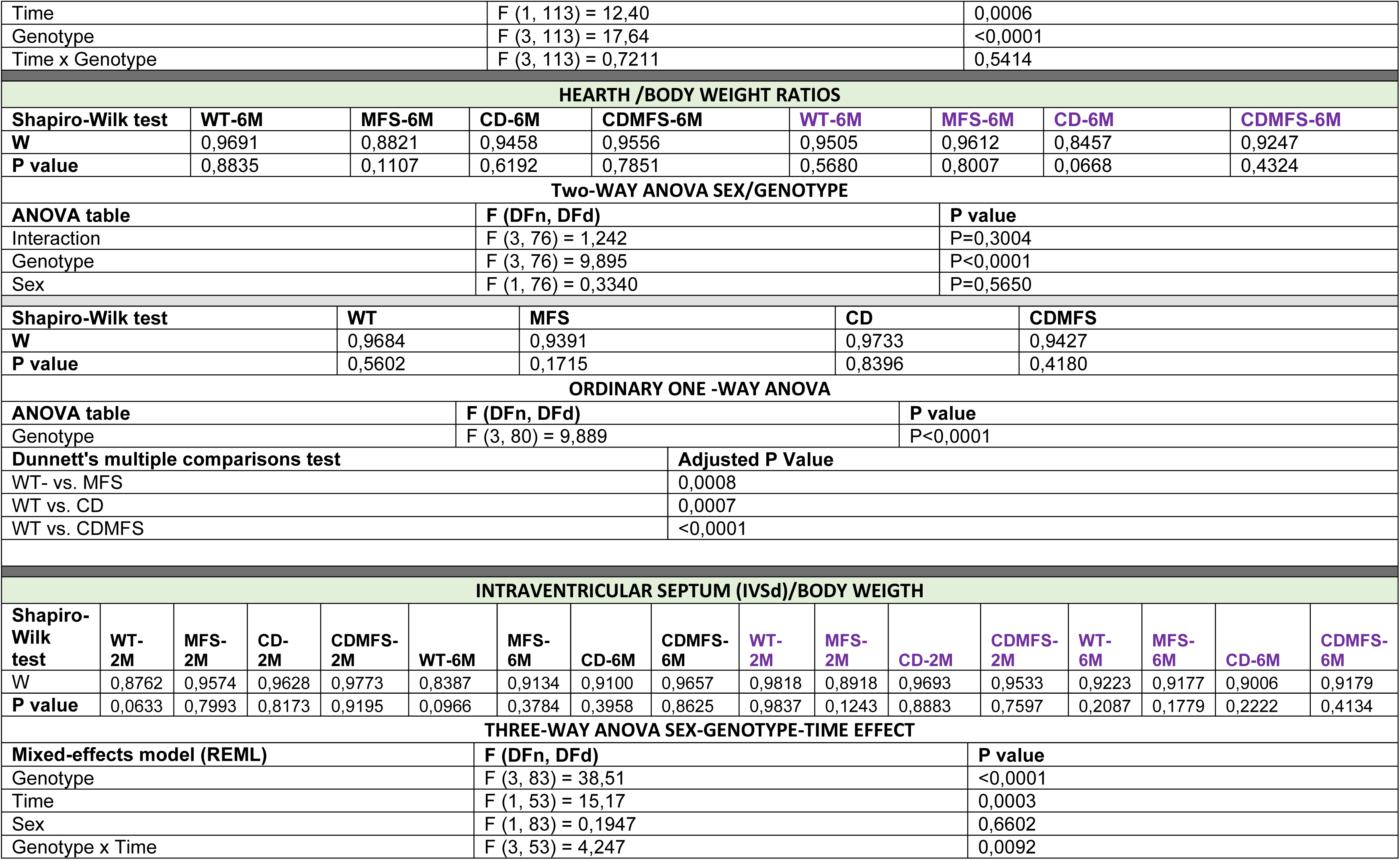
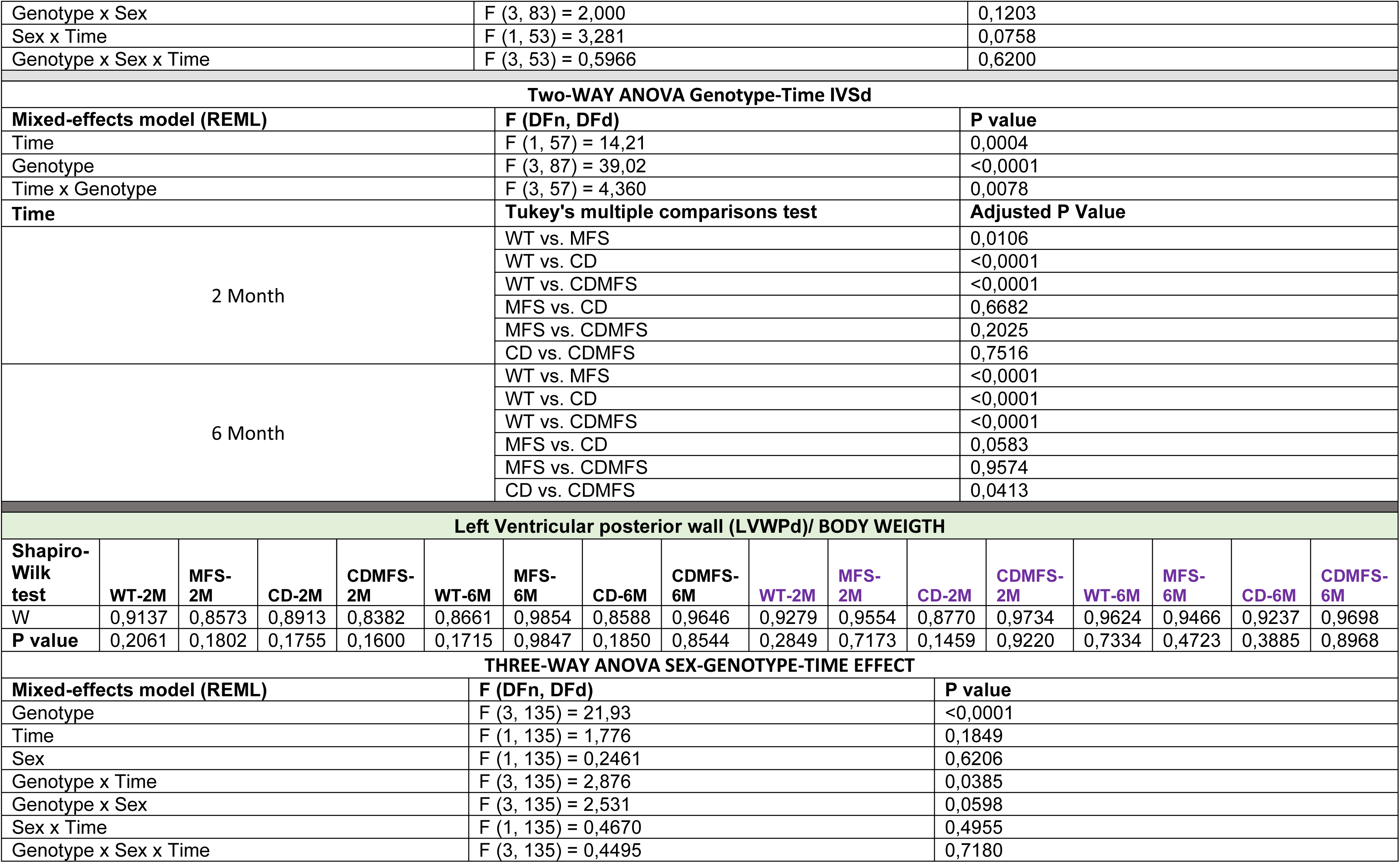
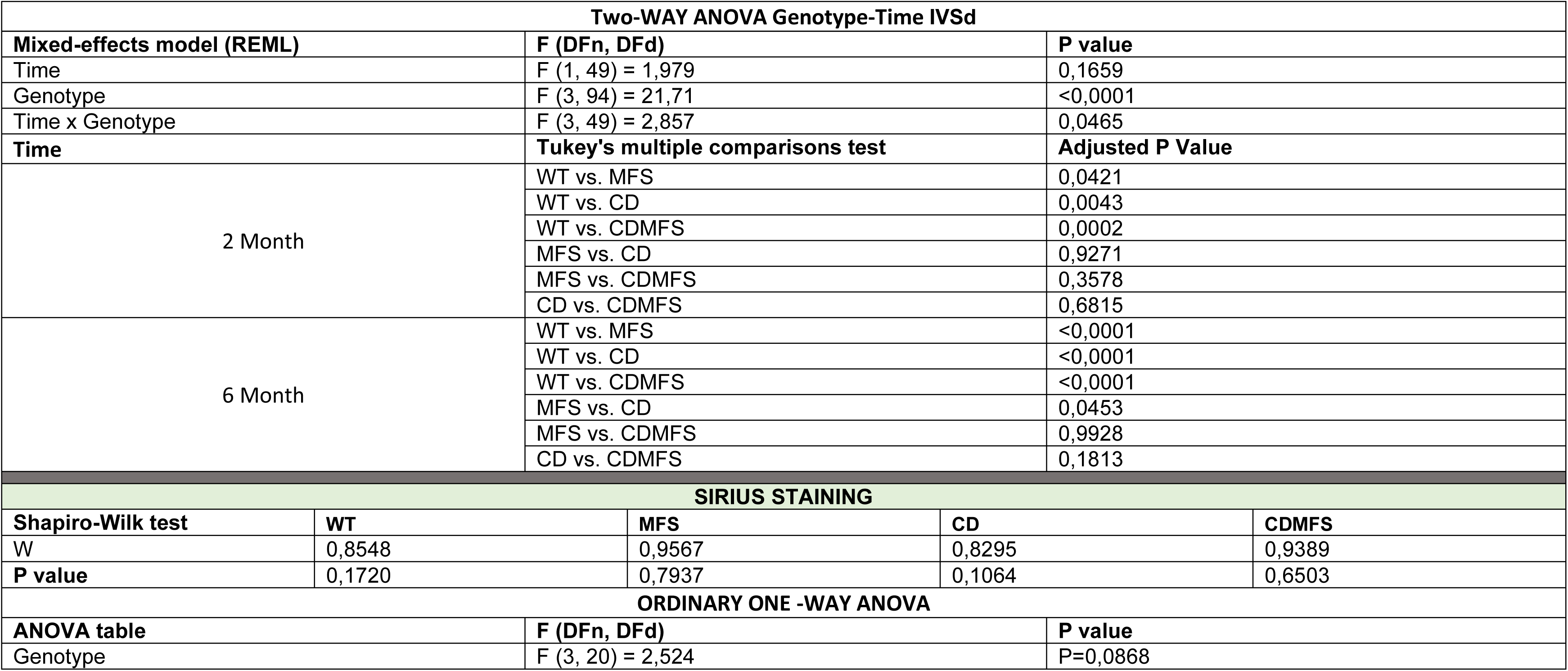
CARDIACS DISFUNCTION.

**TABLE S8:** REDOX STRESS MARKERS.

| OXPHOS COMPLEXES |  |  |  |  |  |  |  |  |
| --- | --- | --- | --- | --- | --- | --- | --- | --- |
| Kruskal-Wallis test |  |  |  |  |  |  |  |  |
| Kruskal-Wallis statistic |  |  | 21.37 |  |  |  |  |  |
| Number of groups |  |  | 8 |  |  |  |  |  |
| P value |  |  | 0,0033 |  |  |  |  |  |
| Dunn's multiple comparisons test |  |  | Adjusted P Value |  |  |  |  |  |
| WT-Males vs. WT-Female |  |  | >0,9999 |  |  |  |  |  |
| MFS-Male vs. MFS-Female |  |  | >0,9999 |  |  |  |  |  |
| CD-Male vs. CD-Female |  |  | >0,9999 |  |  |  |  |  |
| CDMFS-Male vs. CDMF-Female |  |  | >0,9999 |  |  |  |  |  |
| Shapiro-Wilk test | WT | MFS | CD | CDMFS |  |  |  |  |
| W | 0,9245 | 0,9345 | 0,9473 | 0,8767 |  |  |  |  |
| P value | 0,4304 | 0,4932 | 0,6838 | 0,3247 |  |  |  |  |
| ORDINARY ONE -WAY ANOVA |  |  |  |  |  |  |  |  |
| ANOVA table |  | F (DFn, DFd) |  | P value |  |  |  |  |
| Genotype |  | F (3, 27) = 26,15 |  | P<0,0001 |  |  |  |  |
| Dunnett's multiple comparisons test |  |  | Adjusted P Value |  |  |  |  |  |
| WT- vs. MFS |  |  | 0,0188 |  |  |  |  |  |
| WT vs. CD |  |  | <0,0001 |  |  |  |  |  |
| WT vs. CDMFS |  |  | 0,0127 |  |  |  |  |  |
| DHE LEVELS |  |  |  |  |  |  |  |  |
| Shapiro-Wilk test | WT | MFS | CD | CDMFS | WT | MFS | CD | CDMFS |
| W | 0,9278 | 0,9568 | 0,9606 | 0,8999 | 0,9234 | 0,9273 | 0,9444 | 0,9106 |
| P value | 0,5326 | 0,7947 | 0,8240 | 0,3302 | 0,5524 | 0,5282 | 0,6975 | 0,4002 |
| Two-WAY ANOVA SEX/GENOTYPE |  |  |  |  |  |  |  |  |
| ANOVA table |  |  | F (DFn, DFd) |  |  | P value |  |  |
| Interaction |  |  | F (3, 41) = 0,8410 |  |  | P=0,4793 |  |  |
| Sex |  |  | F (1, 41) = 0,06844 |  |  | P=0,7949 |  |  |
| Genotype |  |  | F (3, 41) = 11,83 |  |  | P<0,0001 |  |  |
| Shapiro-Wilk test | WT |  | MFS |  | CD |  | CDMFS |  |
| W | 0,9118 |  | 0,9333 |  | 0,9636 |  | 0,9497 |  |
| P value | 0,2253 |  | 0,3763 |  | 0,8155 |  | 0,5940 |  |

| ORDINARY ONE -WAY ANOVA |  |  |  |  |  |  |  |  |
| --- | --- | --- | --- | --- | --- | --- | --- | --- |
| ANOVA table |  |  | F (DFn, DFd) |  |  | P value |  |  |
| Genotype |  |  | F (3, 45) = 12,56 |  |  | P<0,0001 |  |  |
| Dunnett's multiple comparisons test |  |  |  |  | Adjusted P Value |  |  |  |
| WT- vs. MFS |  |  |  |  | >0,9999 |  |  |  |
| WT vs. CD |  |  |  |  | 0,0001 |  |  |  |
| WT vs. CDMFS |  |  |  |  | 0,6024 |  |  |  |
| 3'-NT LEVELS |  |  |  |  |  |  |  |  |
| Shapiro-Wilk test | WT | MFS | CD | CDMFS | WT | MFS | CD | CDMFS |
| W | 0,9127 | 0,8004 | 0,8810 | 0,7242 | 0,8661 | 0,9604 | 0,7899 | 0,9116 |
| P value | 0,4151 | 0,0593 | 0,2738 | 0,0110 | 0,2111 | 0,8217 | 0,0669 | 0,4473 |
| Kruskal-Wallis test |  |  |  |  |  |  |  |  |
| Kruskal-Wallis statistic |  |  |  |  | 19.45 |  |  |  |
| Number of groups |  |  |  |  | 8 |  |  |  |
| P value |  |  |  |  | 0,0069 |  |  |  |
| Dunn's multiple comparisons test |  |  |  |  | Adjusted P Value |  |  |  |
| WT-Males vs. WT-Female |  |  |  |  | >0,9999 |  |  |  |
| MFS-Male vs. MFS-Female |  |  |  |  | >0,9999 |  |  |  |
| CD-Male vs. CD-Female |  |  |  |  | >0,9999 |  |  |  |
| CDMFS-Male vs. CDMF-Female |  |  |  |  | >0,9999 |  |  |  |
| Shapiro-Wilk test |  | WT |  | MFS |  | CD |  | CDMFS |
| W |  | 0,9536 |  | 0,9171 |  | 0,7940 |  | 0,7933 |
| P value |  | 0,6532 |  | 0,2289 |  | 0,0174 |  | 0,0077 |
| Kruskal-Wallis test |  |  |  |  |  |  |  |  |
| Kruskal-Wallis statistic |  |  |  |  | 18.09 |  |  |  |
| Number of groups |  |  |  |  | 4 |  |  |  |
| P value |  |  |  |  | 0,0004 |  |  |  |
| Dunn's multiple comparisons test |  |  |  |  | Adjusted P Value |  |  |  |
| WT- vs. MFS |  |  |  |  | 0,0344 |  |  |  |
| WT vs. CD |  |  |  |  | 0,0001 |  |  |  |
| WT vs. CDMFS |  |  |  |  | 0.5956 |  |  |  |

**TABLE S9:** MMP2 Expression.

| MMP2 Expression |  |  |  |  |
| --- | --- | --- | --- | --- |
| Kruskal-Wallis test |  |  |  |  |
| Kruskal-Wallis statistic |  |  | 23.84 |  |
| Number of groups |  |  | 8 |  |
| P value |  |  | 0.0012 |  |
| Dunn's multiple comparisons test |  |  | Adjusted P Value |  |
| WT-Males vs. WT-Female |  |  | >0,9999 |  |
| MFS-Male vs. MFS-Female |  |  | >0,9999 |  |
| CD-Male vs. CD-Female |  |  | >0,9999 |  |
| CDMFS-Male vs. CDMF-Female |  |  | >0,9999 |  |
| Shapiro-Wilk test | WT | MFS | CD | CDMFS |
| W | 0,9340 | 0,8873 | 0,9278 | 0,7829 |
| P value | 0,4523 | 0,1871 | 0,4604 | 0,0583 |
| ORDINARY ONE -WAY ANOVA |  |  |  |  |
| ANOVA table |  | F (DFn, DFd) | P value |  |
| Genotype |  | F (3, 30) = 11,97 | P<0,0001 |  |
| Dunnett's multiple comparisons test |  |  | Adjusted P Value |  |
| WT- vs. MFS |  |  | 0,0006 |  |
| WT vs. CD |  |  | 0,4095 |  |
| WT vs. CDMFS |  |  | 0.0312 |  |

**TABLE S10.** Summary of main results in the CDMFS mouse model.

|  |  | **CD-like phenotype**<br>**(CD > MFS)** | **MFS-like phenotype**<br>**(MFS > CD)** | **CD =/~ MFS phenotype** | **CD >< MFS**<br>**(tug-of-force)**<br>**phenotype** |
| --- | --- | --- | --- | --- | --- |
| **Brain-based**<br>**outcomes** |  | Marble buried test<br>(--)<br>Fig. 1A, B | ----- | ----- | ----- |
|  |  | Brain weight<br>(--)<br>Fig. 1C | ----- | ----- | ----- |
| **Cardiovascular** | **Aortic outcomes** | Total aortic area<br>(the media + lumen)<br>(--)<br>Fig. 3A | Aortic root diameter<br>(++)<br>Fig. 2 | Tunica media only<br>(+++)<br>Fig. 3C | Tunica media cell<br>density<br>(±)<br>Fig. 3D |
|  |  | Blood pressure<br>(++)<br>(Fig. 5) | Elastic fiber undulation<br>(++)<br>Fig. 4C | ----- | Elastic fibres breaks (+)<br>Figs. 4A, B |
|  | **Cardiac outcomes** | ----- | Mitochondria OXPHOS<br>(-)<br>Fig. 7A | Ejection fraction (EF)<br>(-)<br>Fig. 6B, C, D | MMP-2<br>(+)<br>Fig. 8 |
|  |  | ----- | DHE (redox stress)<br>(=)<br>Fig. 7B | Heart/body weight<br>(+++)<br>Fig. 6A | 3-NT (redox stress)<br>(=)<br>Fig. 7C |

Meaning of symbols: (+) augmented; (-) reduced; (=/~) no/similar; (±), intermediate.
Each outcome is referenced to its respective figure in the text for details.

